# B7-H4 Binds Galectin-9 Glycosylation-Dependently and Attenuates Galectin-9-Mediated CD28/AKT Activation and T Cell Death

**DOI:** 10.1101/2025.07.31.666261

**Authors:** Ravear Zhiqiang Wang, Fang Yang, Jianhua Sui

## Abstract

B7-H4, a member of the B7 family, is broadly expressed on various cancer cells and has been implicated in negative immune regulation, particularly in suppressing anti-tumor immunity. However, the receptor for B7-H4 and the molecular mechanisms underlying its immunoinhibitory effects remain poorly understood. In this study, using peritoneal immune cells from mice adoptively transferred with OVA-expressing tumor cells and OVA-specific OT-1 T cells, we identified Galectin-9 (Gal-9) as a binding partner for B7-H4 and elucidated its role in modulating T cell responses through this interaction. We demonstrated that glycosylation in the IgC domain of B7-H4 is required for its binding to Gal-9; while the N-terminal carbohydrate recognition domain (N-CRD) of Gal-9, including the R65 residue in the N-CRD, is essential for this interaction. Additionally, we found that other B7 family members (B7.1, B7.2, B7-H2, and B7-DC) and immune cell surface receptors (CD28, 2B4, CD226, and SLAMF1) also bind to Gal-9 at comparable levels to B7-H4 and T cell immunoglobulin mucin receptor 3 (TIM-3). *In vitro* functional assays revealed that B7-H4 inhibits Gal-9-induced activation of CD28 downstream signaling and reduces Gal-9-mediated T cell death. *In vivo*, Gal-9 deficiency in mice resulted in a significant reduction in the proportion of splenic CD4^+^ T cells, whereas B7-H4 deficiency exhibited no observable phenotype. Furthermore, B7-H4 and Gal-9 double-knockout mice displayed no additional phenotype differences compared to Gal-9 single-knockout mice. Notably, tumor growth following tumor cell challenge was unaffected in all three knockout models (Gal-9 single-, B7-H4 single-, or double-knockout). Collectively, these findings suggest that B7-H4, other B7 family members, Gal-9, and T cell surface immune receptors form a complex regulatory network that modulates T cell activity and anti-tumor responses, although no single member exerts a major effect. This study provides a detailed molecular characterization of the interaction between B7-H4 and Gal-9 and identifies other previously unknown Gal-9 binding partners, offering valuable insights into the intricate regulatory network involving these molecules.

## Introduction

B7-H4 (B7 homolog 4, also known as B7S1, B7x, or VTCN1) is an orphan immune checkpoint ligand belonging to the B7 family. It has been proposed that B7-H4 negatively regulates T cell immune response through multiple mechanisms, including the suppression of T cell proliferation in response to antigen stimulation (1), the reduction of key cytokines, such as IL-2 and IFN-γ (2), the induction of T cell anergy (3), and the protection of normal tissues from T cell-mediated destruction (4). Moreover, B7-H4 is frequently overexpressed in various cancer cells, where it acts as a negative regulator of anti-tumor T cell immunity (5–8). Beyond its expression on tumor cells, cell-surface B7-H4 is also expressed on tumor-associated macrophages, where it further suppresses T cell-mediated immune responses (9). These findings underscore the role of B7-H4 in regulating immune responses in both physiological and pathological contexts. However, the identity of its receptor and the precise molecular mechanisms through which B7-H4 exerts its immunoinhibitory activity remain unclear (10).

A previous study reported that B7-H4 interacts with activated T lymphocytes from wild type (WT) mice, but not with T cells from B and T lymphocyte attenuator (BTLA)-deficient mice (11), indirectly suggesting that BTLA could be a potential receptor for B7-H4. However, subsequent studies have shown that BTLA is unlikely to function as a receptor for B7-H4, and herpesvirus entry mediator (HVEM) has been identified as the ligand for BTLA (12, 13). Additionally, another study suggests that B7-H4 associates with the soluble semaphoring 3a (Sema3a), which acts as a functional bridge to stimulate a Nrp-1/Plexin A4 heterodimer, forming an immunoregulatory receptor complex that modulates the activity of regulatory CD4^+^ T cells (14). Nevertheless, Sema3a itself does not serve as a direct receptor for B7-H4. Intriguingly, other research indicates that tumor-infiltrating CD8^+^ T cells may express a putative receptor for B7-H4, as antigen-presenting cells (APCs) expressing B7-H4 suppress the effector functions of CD8^+^ T cells (15). These findings highlight the complexity of B7-H4-dependent immune regulation and underscore the need for further investigation.

Herein, we identify Galectin-9 (Gal-9) as a previously unknown binding partner for B7-H4 using an immunoprecipitation (IP) coupled with mass spectrometry (MS) approach, where recombinant B7-H4 protein served as a bait, while peritoneal immune cells—harvested from mice adoptively transferred with EG7 tumor cells (an OVA-expressing derivative of the mouse T cell lymphoblast EL4 cell line) and OVA-specific T cells (OT-1 T cells)—were used as target cells. Further characterization of the interaction between B7-H4 and Gal-9 revealed that glycosylation in the IgC domain of B7-H4 is required for its binding to Gal-9; while the N-terminal carbohydrate recognition domain (N-CRD) and the R65 residue within the N-CRD of Gal-9 are essential for this interaction. Notably, we also discovered that Gal-9 binds several previously unrecognized immune receptors on T cells, including CD28, and other B7 family members. Soluble glycosylated B7-H4 attenuates Gal-9’s dual activities: the stimulation of CD28/AKT signaling and the induction of T cell death. Furthermore, a comparison of the splenic immune cell compositions revealed that the pattern and extent of the increased proportion of CD4^+^ T cells in Gal-9 knockout (KO) mice was similar to that in B7-H4 and Gal-9 double knockout (DKO) (B7-H4/Gal-9 DKO) mice, whereas B7-H4 KO mice showed no such increase, indicating that B7-H4 KO alone is insufficient to alter Gal-9’s activity in this context *in vivo*, consistent with the observation that multiple B7 family members and immune receptors on T cells interact with Gal-9. Our findings suggest that B7-H4, other B7 family members, and T cell surface immune receptors interact with Gal-9 within a complex regulatory network that finely modulates T cell activity through multiple overlapping pathways.

## Results

### Identification of Galectin-9 (Gal-9) as a B7-H4-binding protein

To elucidate the molecular mechanism underlying B7-H4’s negative immune-regulatory function, we first aimed to identify its potential binding partner(s). Preliminary evidence suggests that B7-H4 interacts with molecules expressed on activated—but not resting—T cells (15). However, using recombinant human B7-H4 protein (hB7-H4-Fc, consisting of the human B7-H4 ectodomain, including both the IgV and IgC domains, fused to the mouse IgG2a Fc region), we found that the B7-H4 did not bind to activated primary human T cells (**Fig. S1A**). Similarly, using recombinant mouse B7-H4 protein (mB7-H4-Fc, consisting of the mB7-H4 ectodomain, including both the IgV and IgC domains, fused to the human IgG1 Fc region), we found that mB7-H4 did not bind to naïve or antigen (OVA)-activated OT-1 T cells (**Fig. S1B**).

Since mouse B7-H4 has been shown to be expressed on mouse peritoneal macrophages (3), we hypothesized that a putative mB7-H4 receptor may exist on immune cells within the mouse peritoneal cavity contributing to its immune-regulating function. To test this hypothesis, we first assessed the binding of mB7-H4-Fc to peritoneal immune cells from C57BL/6J mice (CD45.1^+^, CD45.2^-^) using flow cytometry. We found that only 1.57% of the CD45.1^+^ peritoneal immune cells were positive for mB7-H4-Fc binding (**Fig. 1A**). In contrast, when analyzing peritoneal immune cells isolated from the same strain of mice one day after the peritoneally adoptive transfer of EG7 tumor cells and OVA-specific OT-1 T cells (CD45.1^-^, CD45.2^+^), we observed that 15.5% of the total CD45.1^+^ host peritoneal immune cells were positive for mB7-H4-Fc binding (**Fig. 1A**); the CD45.1^-^ OT-1 T cells in the peritoneal cavity were again found negative for mB7-H4-Fc binding (**Fig. S1C**). These findings suggest that a putative mB7-H4 binding protein may be expressed or upregulated on the peritoneal immune cells in mice following the peritoneal adoptive transfer with EG7 tumor cells and OT-1 T cells.

**Fig. 1.**
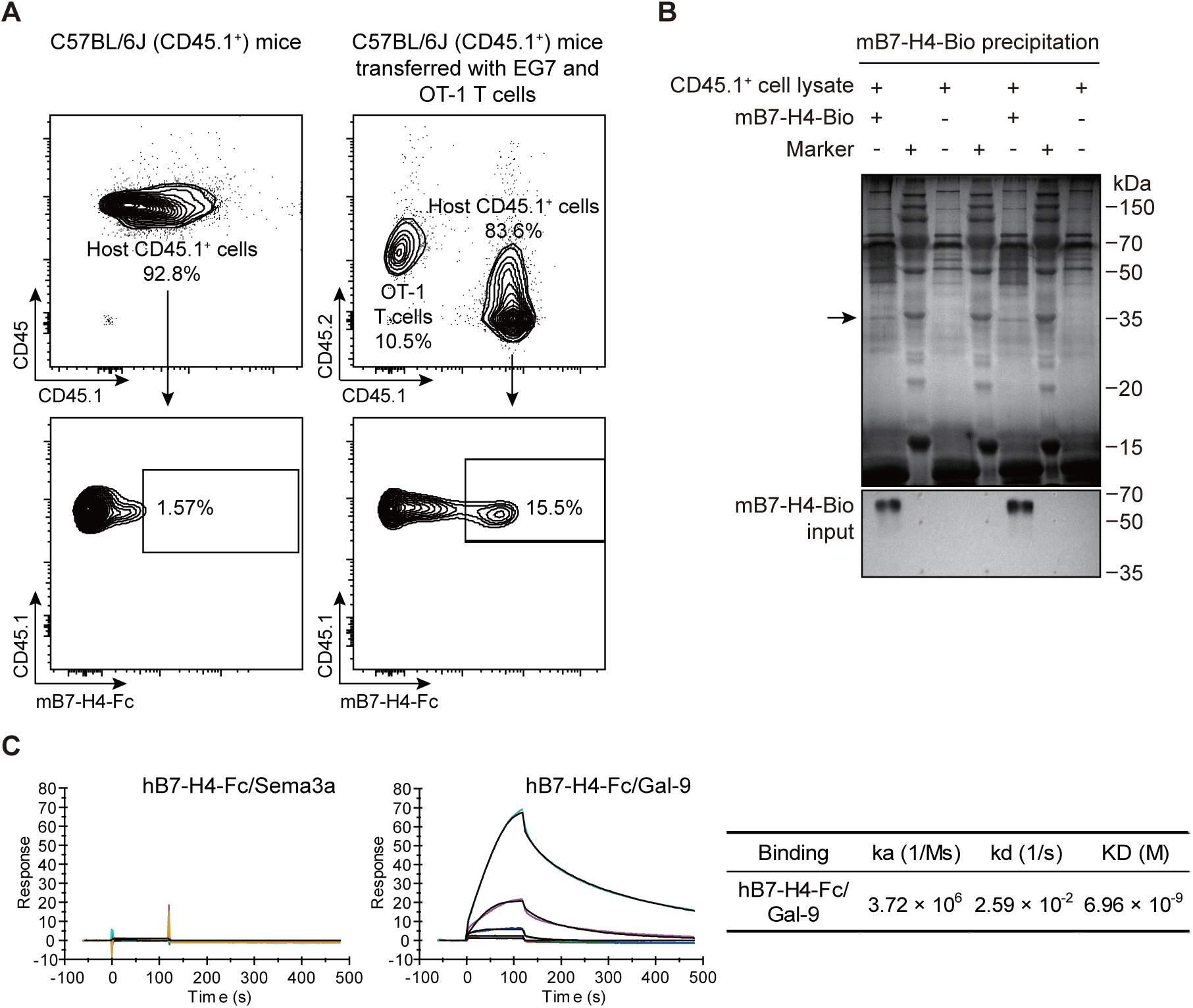
Identification of B7-H4’s binding partner, Gal-9. **(A)** Flow cytometry analysis of mB7-H4-Fc binding to host CD45.1^+^ cells in peritoneal immune cells. Peritoneal immune cells were isolated from C57BL/6J mice (CD45.1^+^, CD45.2^-^) mice transferred with or without EG7 tumor cells and OT-1 T cells (see Methods). Rectangle gates represent the proportion of mB7-H4-Fc positive binding within the parent gate of host CD45.1^+^ cells. **(B)** Immunoprecipitation (IP) of peritoneal CD45.1⁺ cell lysates using biotinylated mB7-H4 (mB7-H4-Bio). Streptavidin beads captured mB7-H4-Bio-precipitated proteins were separated by SDS-PAGE and visualized by silver staining. The arrow indicates a specific ∼35 kDa band precipitated by mB7-H4-Bio. The input mB7-H4-Bio protein was confirmed by western blotting with HRP-Streptavidin. **(C)** SPR binding kinetics between hB7-H4-Fc and Gal-9. The Sema3a showed no binding to hB7-H4-Fc, while Gal-9 exhibited specific binding. The binding kinetic parameters (ka, kd, KD) are shown on the right.

To identify candidate binding proteins for mB7-H4, we next employed an immunoprecipitation (IP) coupled with mass spectrometry (MS) method. A biotinylated mB7-H4 recombinant protein (mB7-H4-Bio, see Methods) and M-280 Streptavidin magnetic beads were used to precipitate mB7-H4-Bio-binding proteins from the cell lysates of CD45.1^+^ peritoneal immune cells from the adoptive-transfer-model mice described above. Cell lysates from EG7 and 4T1 (a mouse breast cancer cell line) tumor cells were used as controls. The precipitated proteins were separated by SDS-PAGE electrophoresis and visualized by silver staining. A protein band with an apparent molecular weight of approximately 35 kDa was observed only when both mB7-H4-Bio and CD45.1^+^ peritoneal immune cell lysates were present; whereas this band was not detected when each component was tested alone or under other conditions, including when both mB7-H4-Bio and 4T1 or EG7 cell lysates were present (**Fig. 1B and Fig. S1D**). Next, this ∼35 kDa band was excised from the gel, digested with trypsin, and subjected to MS analysis using LTQ-Orbitrap Velos (Thermo Fisher Scientific). The MS analysis identified mouse Galectin-9 (mGal-9) exhibited the highest abundance among all candidate binding proteins, with seven distinct mGal-9-matching peptides covering 26% of its protein sequence (**Fig. S1E**). Notably, the theoretical molecular weight of mGal-9 (322 amino acids, 36.5 kDa), closely corresponds to the observed ∼35 kDa band of interest. These results indicate that mGal-9 is a potential binding protein for mB7-H4.

To investigate whether the binding between mB7-H4 and mGal-9 is conserved in humans, we examined the binding of human B7-H4 to human Gal-9. Using surface plasmon resonance (SPR) (Biacore T100 system, GE Healthcare), we tested hB7-H4-Fc binding to a purified human Gal-9 protein (R&D systems). The result showed that hB7-H4-Fc specifically bound to the Gal-9 protein, but not to the purified human Sema3a protein (also served as a negative control; **Fig. 1C**). These findings demonstrate the binding of B7-H4 and Gal-9 is conserved across species in both mice and humans.

### Binding of B7-H4 to Gal-9 is primarily glycan-mediated, involving B7-H4 ectodomain’s N-linked glycan, Gal-9’s N-CRD, and the R65 residue within the N-CRD

To further characterize the binding between B7-H4 and Gal-9, we investigated which Ig-like ectodomain of human B7-H4 mediates binding to human Gal-9. Two human B7-H4 ectodomain truncation variant proteins, the IgV-only variant (hB7-H4 IgV-Fc) and the IgC-only variant (hB7-H4 IgC-Fc), were produced similarly as the aforementioned hB7-H4-Fc (containing both IgV and IgC domains). SPR analysis revealed that hB7-H4 IgC-Fc bound to Gal-9 protein at levels comparable to hB7-H4-Fc, whereas the hB7-H4 IgV-Fc exhibited a markedly reduced binding (**Fig. 2A**). Consistently, flow cytometry confirmed that hB7-H4 IgV-Fc showed reduced binding to 293T cells transiently expressing human Gal-9 (293T-Gal-9 cells), compared to both hB7-H4-Fc and hB7-H4 IgC-Fc (**Fig. 2B**). Taken together, these data suggest that the IgC domain of B7-H4 is essential for its binding to Gal-9.

**Fig. 2.**
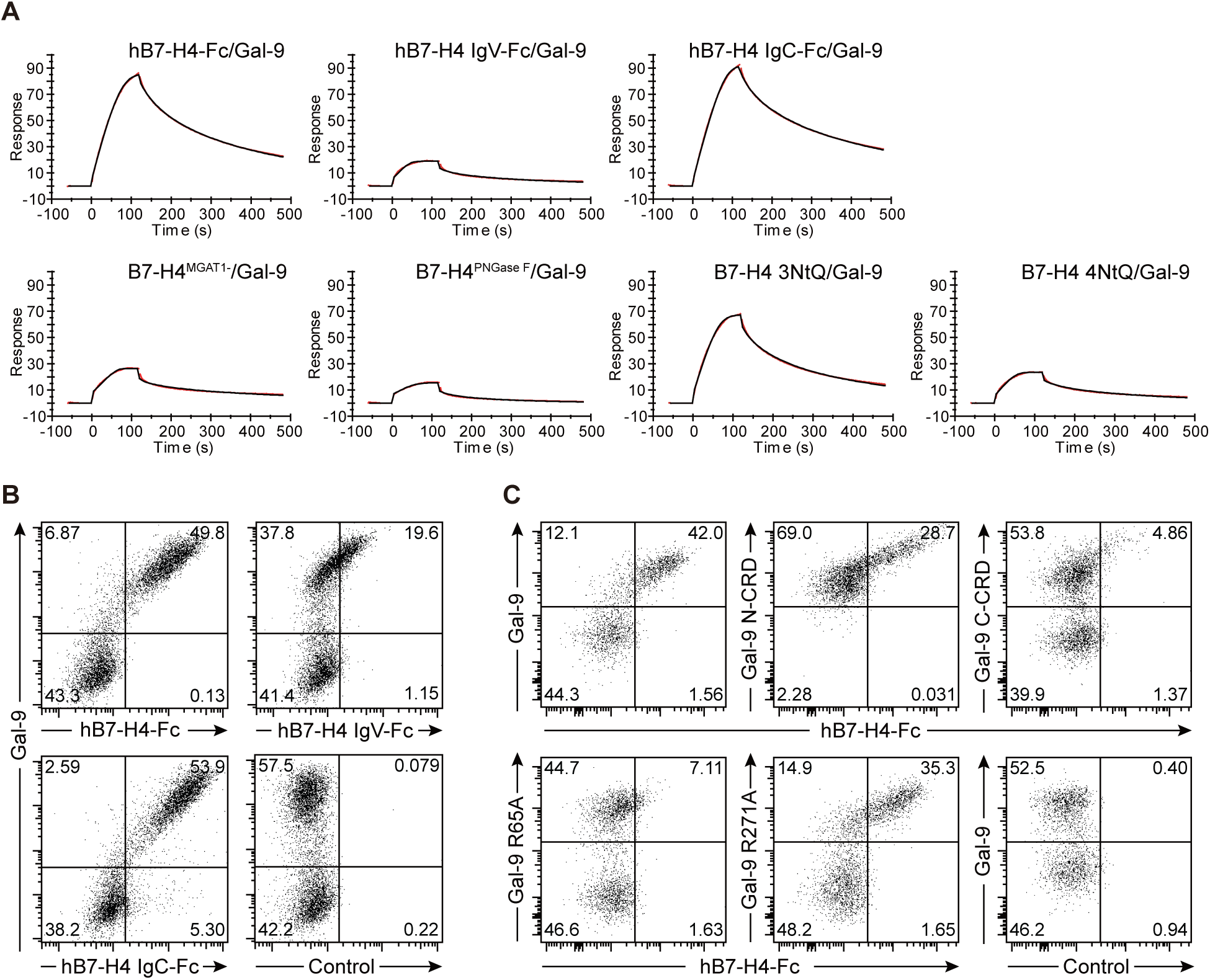
The R65 residue in the Gal-9 N-CRD mediates binding to glycosylated B7-H4 IgC domain. **(A)** The IgC domain and glycosylation of B7-H4 are essential for its binding to Gal-9 protein. SPR binding kinetics are shown for hB7-H4-Fc and its various variants, including truncation variants (hB7-H4 IgV-Fc and hB7-H4 IgC- Fc), de-glycosylated forms (B7-H4^MGAT1-^ and B7-H4^PNGase^ ^F^), and mutants (B7-H4 3NtQ and B7-H4 4NtQ), in their binding to Gal-9 protein. **(B)** The IgC domain of B7-H4 is essential for its binding to Gal-9-expressing cells. hB7- H4-Fc and its truncation variants were analyzed for binding to Gal-9-EGFP-expressing 293T cells by flow cytometry. **(C)** The R65 residue in the N-CRD of Gal-9 is required for its interaction with B7-H4. hB7-H4-Fc was examined for binding to Gal-9-EGFP-expressing 293T cells, Gal-9 truncation variants (Gal-9 N-CRD and Gal-9 C-CRD), or Gal-9 single-site mutants (Gal-9 R65A and Gal-9 R271A)-EGFP-expressing 293T cells by flow cytometry.

Given that Gal-9 is a lectin with carbohydrate recognition capabilities (16) and B7-H4 is a glycoprotein with seven potential N-linked glycosylation sites located across its IgV and IgC domains (1, 5) (**Fig. S2A**), the binding between B7-H4 and Gal-9 likely occurs in a glycan-mediated manner. It has also been reported that B7-H4’s glycosylation plays a crucial role in its stability and function (17). To assess the role of B7-H4’s glycosylation in its binding with Gal-9, hB7-H4-Fc protein were de-glycosylated either by expressing it in N-acetylglucosaminyltransferase I (MGAT1) KO 293F cells, which lack the ability to synthesize complex N-linked glycans (18), and naming it B7-H4^MGAT1-^, or by treating purified hB7-H4-Fc with PNGase F to remove N-linked glycans (B7-H4^PNGase^ ^F^). Both the de-glycosylated proteins, B7-H4^MGAT1-^ and B7-H4^PNGase^ ^F^, exhibited reduced binding to human Gal-9 protein compared to hB7-H4-Fc protein with glycosylation (**Fig. 2A**), indicating the critical role of B7-H4’s glycosylation in mediating its binding with Gal-9.

To further investigate the specific glycosylation site involved, we generated B7-H4 mutants by substituting asparagine with glutamine at the predicted N-linked glycosylation sites using hB7-H4-Fc protein as the backbone to produce seven single-site mutants: N112Q in the IgV domain, and six in the IgC domain (N160Q, N190Q, N196Q, N205Q, N216Q, and N220Q), and two multi-site mutants: B7-H4 3NtQ (N205Q, N216Q, and N220Q) and B7-H4 4NtQ (N112Q, N160Q, N190Q, and N196Q) (**Fig. S2A–B**). SPR analysis showed that all seven single-site mutants (**Fig. S2C**) and the multi-site mutant B7-H4 3NtQ exhibited minimal changes in Gal-9 binding compared to WT hB7-H4-Fc; whereas the B7-H4 4NtQ mutant showed weaker Gal-9 binding than WT hB7-H4-Fc (**Fig. 2A**). These results demonstrate that glycosylation at N160, N190, and N196 in the IgC domain of human B7-H4 ectodomain are critical for its binding to human Gal-9, consistent with the observation that the IgC domain is essential for this interaction.

Human Gal-9 consists of two CRDs, N-terminal CRD (N-CRD) and C-terminal CRD (C-CRD). To identify which CRD of Gal-9 is involved in binding with human B7-H4, we transiently overexpressed each of the CRD domain of human Gal-9 (UniProt O00182-1) in 293T cells and generated the N-CRD-only variant (293T-Gal-9 N-CRD) cells and the C-CRD-only variant (293T-Gal-9 C-CRD) cells, similar to that for generating WT 293T-Gal-9 cells described above. The binding of hB7-H4-Fc to these cells was assessed using flow cytometry. We found that hB7-H4-Fc bound to 293T-Gal-9 N-CRD cells at slightly lower levels compared to WT 293T-Gal-9 cells, whereas its binding to 293T-Gal-9 C-CRD cells was markedly weaker (**Fig. 2C**). To further investigate the specific residues involved in the binding, two Gal-9’s loss-of-function single-site mutants (19), R65A in the N-CRD and R271A in the C-CRD (corresponding to R239A in the short isoform of Gal-9 (UniProt O00182-2)) were similarly expressed in 293T cells. Flow cytometry analysis revealed that hB7-H4-Fc bound to 293T-Gal-9 R271A cells at levels comparable to 293T-Gal-9 cells, whereas its binding to 293T-Gal-9 R65A cells was markedly reduced (**Fig. 2C**). Taken together, these results demonstrate that the binding between human B7-H4 and human Gal-9 is mediated by the glycosylated B7-H4 ectodomain, particularly the IgC domain, and the N-CRD domain of Gal-9, with the R65 residue in the N-CRD serving as a critical determinant of this binding.

### B7-H4 binding to Gal-9 attenuates Gal-9-induced T cell death

Gal-9 is recognized as an immune-regulator with diverse but not yet fully understood mechanisms. It plays a dual role in immunoregulation, capable of either stimulating or suppressing the immune response (20). Its activity on T cells is believed to depend on whether it is localized intracellularly or extracellularly (21, 22). While Gal-9 is predominantly localized within intracellular compartments in CD4^+^ and CD8^+^ T cells, it translocates to the T cell membrane upon activation. Notably, exogenous soluble Gal-9 is recognized for its role in inducing T cell apoptosis (16, 23, 24), a process initially believed to be mediated by TIM-3 expressed on T cells (16). However, subsequent studies revealed that TIM-3 is not the sole mediator of this mechanism (25, 26). This prompted us to first investigate whether B7-H4 participates in Gal-9-induced T cell apoptosis.

We first sought out to confirm that Gal-9 induces T cell death in primary human T cells. To this end, human peripheral blood mononuclear cells (PBMCs) were incubated with Gal-9 for 0.5 hours (h), and the cell viability was analyzed using flow cytometry. Compared to untreated control, Gal-9 treatment significantly reduced cell viability: total PBMCs decreased from ∼ 80% to ∼ 48%, and CD3^+^ T cells declined from ∼ 62% to ∼ 48% (**Fig. S3A–B**). In contrast, B cell viability was not markedly affected by Gal-9 treatment (**Fig. S3B**). Likewise, directly treating primary T cells isolated from PBMCs with Gal-9 for either 0.5 h or 18 h also significantly reduced the viability of T cells (**Fig. S3C–D**).

Having confirmed that Gal-9 induces T cell death, we next tested whether B7-H4 inhibits this effect in primary T cells, Jurkat T cells (a human T lymphoblastoid cell line), and MOLT-4 T cells (a human T lymphoblast cell line). Each of these cell types were incubated with Gal-9 for 0.5 h with or without hB7-H4-Fc or de-glycosylated B7-H4^MGAT1^, followed by cell viability analysis using flow cytometry. We found that hB7-H4-Fc markedly reduced Gal-9-induced T cell death, whereas B7-H4^MGAT1-^ had much less or minimal effect in all the three cell types examined, including primary T cells (**Fig. 3A**), Jurkat T cells (**Fig. 3B**) and MOLT-4 T cells (**Fig. 3C**). Another form of B7-H4, iB7-H4 (immobilized hB7-H4-Fc on Protein A beads), similarly reduced Gal-9-induced T cell death and de-glycosylated iB7-H4^MGAT1-^ had no effect on Gal-9-induced T cell death in MOLT-4 T cells (**Fig. 3D**). These results demonstrate B7-H4 attenuates Gal-9-induced T cell death, and this process is glycosylation-dependent, consistent with its glycosylation-dependent binding to Gal-9.

**Fig. 3.**
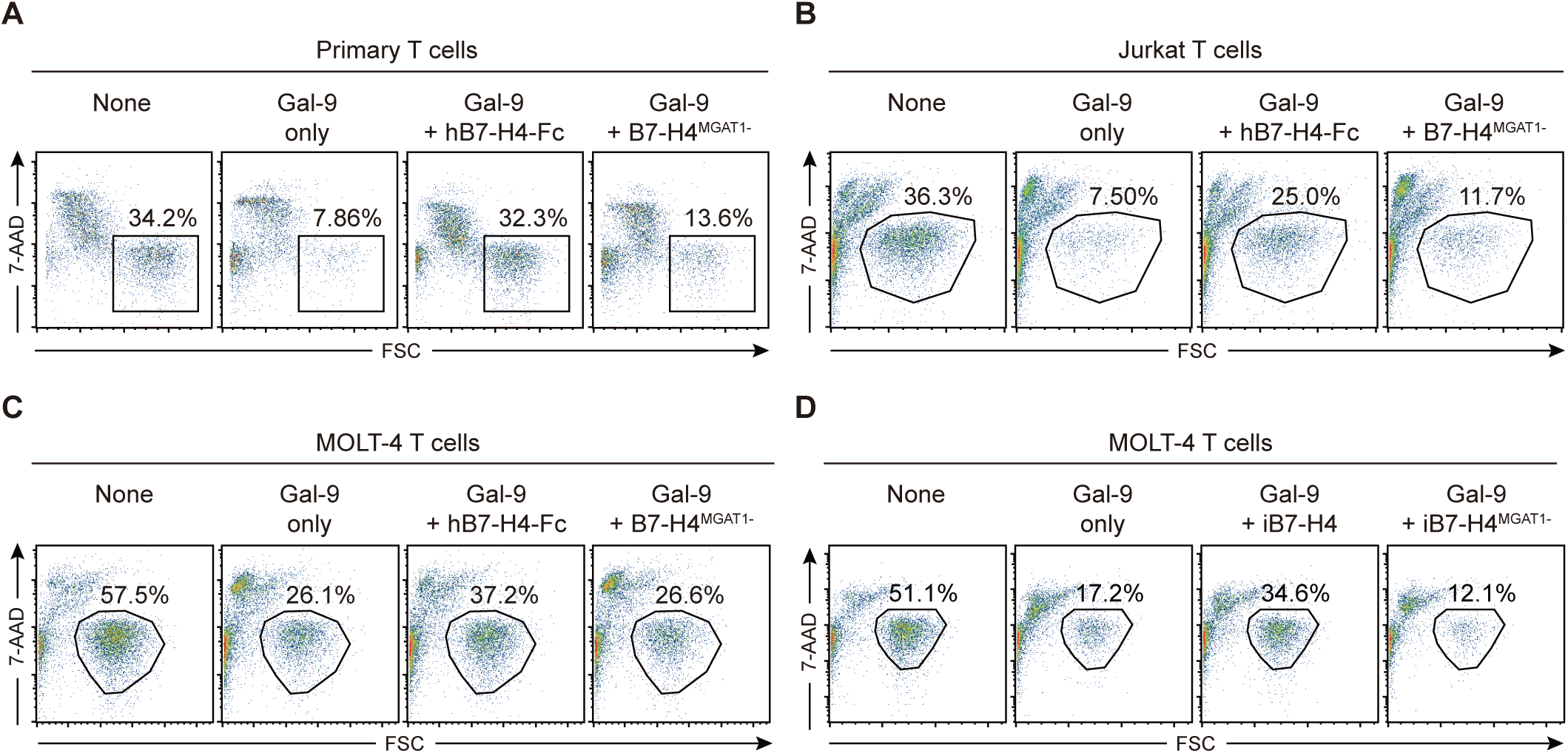
B7-H4 suppresses Gal-9-induced T cell death, and this suppression depends on the glycosylation of B7-H4. (A-C) Flow cytometry analysis of live cells (7-AAD negative) within primary human T cells (A), Jurkat T cells (B), and MOLT-4 T cells (C) after treatment with 8 µg/mL Gal-9 for 30 min in the presence or absence of 25 µg/mL glycosylated hB7-H4-Fc, or de-glycosylated hB7-H4-Fc (B7-H4^MGAT1-^). **(D)** Flow cytometry analysis of the proportion of live T cells (7-AAD negative) within the parent gate of MOLT-4 T cells treated with 6 µg/mL Gal-9 for 30 min in the presence or absence of Protein A-immobilized glycosylated hB7-H4-Fc (iB7-H4) or de-glycosylated hB7-H4-Fc (iB7-H4^MGAT1-^).

Nonetheless, this finding appears contradictory to B7-H4’s known inhibitory immune-regulating role. Since we did not observe direct binding of B7-H4 to activated T cells (**Fig. S1A–B**), we hypothesize that B7-H4’s inhibition of Gal-9-induced T cell death is likely indirect, mediated through competition with Gal-9’s binding partners on T cells. Identifying these other, as-yet-unknown Gal-9 binding partners on T cells may shed light on the potential complexity of B7-H4’s inhibitory immune-regulating mechanism.

### Gal-9 binds to several T cell surface immune receptors, including CD28, and B7-H4’s binding to Gal-9 interferes with Gal-9-induced CD28 activation

To explore this, we first examined the binding of Gal-9 to 11 additional immune receptors primarily expressed on T cells (some are also expressed on NK cells or other immune cells), as well as its two known binding partners, TIM-3 and PD-1. These receptors were produced as recombinant proteins and tested for binding to Gal-9 protein using SPR. Additionally, they were also tested for binding to 293T-Gal-9 cells using flow cytometry. Of the 11 additional receptors tested, CD28, 2B4 (CD244 or SLAMF4), CD226, and SLAMF1 bound to Gal-9 at levels comparable to PD-1, TIM-3, and B7-H4 in SPR and/or flow cytometry assays (**Fig. 4A–B, Fig. S4A–B**). By comparison, the remaining seven immune receptors showed weak or minimal binding to Gal-9 in SPR analysis (**Fig. S4A**).

**Fig. 4.**
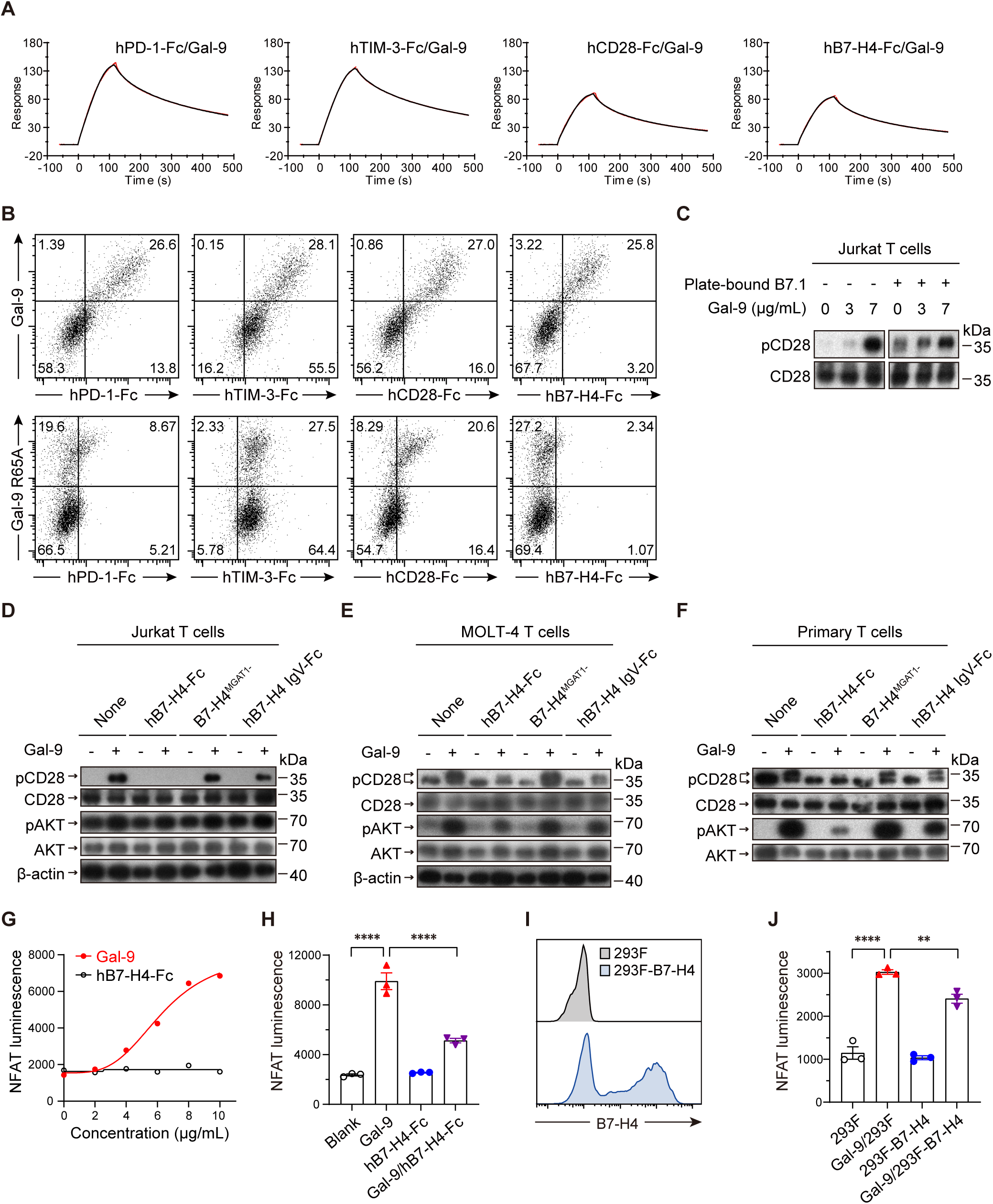
Gal-9 binds to CD28, and the glycosylated B7-H4 attenuates Gal-9-mediated CD28/AKT activation in human T cells. **(A)** SPR analysis of Gal-9 binding kinetics to PD-1, TIM-3, CD28, and B7-H4. Human PD-1, TIM-3, and CD28 were fused to the same mouse IgG2a Fc tags. 1.2 µg/mL Gal-9 was injected over a Protein A/G-coated Biacore chip to assess binding kinetics to each immobilized Fc-fusion protein. **(B)** Flow cytometry analysis of the binding between the four proteins (shown in Panel A) and 293T cells transiently overexpressing either WT Gal-9 or the Gal-9 R65A mutant. **(C)** Gal-9-mediated stimulation of pCD28 signaling activity in Jurkat T cells. The Jurkat cells were treated with or without plate-bound B7.1, and Gal-9’s dose-dependent effects on pCD28 were assessed by western blotting. **(D-F)** B7-H4 modulates pCD28 and pAKT signaling activities in T cells in a glycosylation-dependent manner. Jurkat (D), MOLT-4 (E), and primary T cells (F) were treated with 8 µg/mL Gal-9 in the presence or absence of 25 µg/mL hB7-H4-Fc, hB7-H4 IgV-Fc, or de-glycosylated B7-H4^MGAT1-^. Western blotting was performed to detect pCD28 and pAKT. **(G)** Gal-9 stimulates NFAT signaling. Jurkat-NFAT-Luc reporter cells showed dose-dependent luciferase induction by Gal-9, but not hB7-H4-Fc. **(H)** B7-H4 attenuated Gal-9-induced CD28 signaling. Jurkat-NFAT-Luc reporter cells were treated with 3.3 µg/mL Gal-9, 25 µg/mL hB7-H4-Fc, or both. **(I)** Flow cytometry analysis of B7-H4 expression on 293F cells. B7-H4 expression on 293F cells (293F-B7-H4) was confirmed, the 293F-B7-H4 cells were used in panel J. **(J)** Cell-surface B7-H4 attenuated Gal-9-induced CD28 signaling. Jurkat-NFAT-Luc reporter cells were co-cultured with 293F or 293F-B7-H4 cells in the presence or absence of 3.3 µg/mL Gal-9. One-way ANOVA was used to statistically analyze the mean of NFAT-mediated luminescence signals among groups in Panels H and J. All data are presented as mean ± s.e.m.

Comparative analysis revealed that 2B4, CD28, and SLAMF1, similar to TIM-3, bound to 293T-Gal-9 R65A cells at levels comparable to or only slightly reduced relative to 293T-Gal-9 cells. Conversely, CD226, like PD-1 and B7-H4, showed markedly reduced binding to 293T-Gal-9 R65A cells compared to 293T-Gal-9 cells (**Fig. 4B and Fig. S4B**). These results demonstrate that Gal-9 R65 residue in N-CRD differentially mediates interactions between Gal-9 and its binding partners. This is consistent with established findings that the Gal-9 R65 residue is dispensable for Gal-9’s binding to TIM-3 but plays a role in its binding to PD-1 (19).

Among the four additional Gal-9 binding immune receptors we identified above, CD28 is a pivotal co-stimulatory receptor that potentiates TCR signaling to drive T cell expansion, cytokine secretion, and survival (27). We therefore focused our efforts on analyzing the effect of Gal-9 on T cells through CD28 engagement and whether B7-H4’s binding to Gal-9 indirectly interferes with this effect. As phosphorylation of CD28 (UniProt P10747-1) at tyrosine 191 (pCD28) is essential for PI3K-AKT pathway activation and subsequent T cell proliferation (28, 29), we assessed Gal-9’s effect on CD28 signaling. In Jurkat T cells, Gal-9 induced pCD28 similarly to the plate-bound B7.1, a canonical CD28 agonist. Furthermore, Gal-9 exhibited additive effects with B7.1 in inducing pCD28, as evidenced by the increased phosphorylation levels of CD28 in the presence of plate-bound B7.1 (**Fig. 4C**).

Gal-9 also induced pCD28 in MOLT-4 T cells, as well as in primary human T cells, consistent with observations in Jurkat T cells (**Fig. 4D–F**). Further testing revealed that soluble hB7-H4-Fc inhibited this Gal-9-induced pCD28 in all three cell types, while the two Gal-9 binding-weakened variants, hB7-H4 IgV-Fc and the de-glycosylated B7-H4^MGAT1-^ showed either abolished or markedly reduced inhibition of pCD28 compared to hB7-H4-Fc. Additionally, Gal-9 induced phosphorylation of AKT (UniProt P31749-1) at serine 473 (pAKT) in both MOLT-4 and primary T cells. The pAKT levels were substantially reduced by hB7-H4-Fc treatment, whereas its two variants exhibited noticeably weaker inhibitory effects on pAKT in both cell types, albeit to varying degrees (**Fig. 4E–F**).

Phosphorylated AKT activates downstream NFAT signaling via GSK3β inhibition, upregulating IL-2 expression and promoting T cell proliferation (27, 30, 31). Using Jurkat-NFAT-Luc reporter cells (see Methods) where NFAT drives luciferase expression, we observed that Gal-9 induced NFAT-driven luciferase activity in a dose-dependent manner, while hB7-H4-Fc showed no such effect even at the highest concentration tested (10 μg/mL) **(Fig. 4G**). Additionally, hB7-H4-Fc potently suppressed Gal-9-mediated NFAT activation (**Fig. 4H**); similarly, 293F cells overexpressing cell-surface B7-H4 (293F-B7-H4, **Fig. 4I**) also significantly inhibited Gal-9-mediated NFAT activation (**Fig. 4J**). Collectively, these results demonstrate that Gal-9’s binding with cell-surface CD28 induces CD28 signaling activation (pCD28/pAKT/NFAT) in human T cells; B7-H4 inhibits this effect, regardless of whether B7-H4 is in its soluble or cell-surface form, likely through competitive binding to Gal-9, indirectly interfering with this signaling activation.

### Other B7 family members also bind to Gal-9, and B7-H4 KO alone is insufficient to alter Gal-9’s immune regulatory activity *in vivo*

Given the similarity of B7-H4 to other members in the B7 family and their extensive glycosylation (10), we next examined whether the other six B7 family members that are not expressed on T cells could also bind to Gal-9 in a manner similar to B7-H4. Using SPR and flow cytometry, we found that soluble extracellular domains of four B7 member proteins—B7-H2, B7.1, B7-DC, and B7.2—bound Gal-9 with equal to or greater affinity than hB7-H4-Fc. In contrast, B7-H3 and B7-H1 exhibited weaker binding activity (**Fig. S5A–B**). These findings suggest that B7 family members, Gal-9, and other immune cell surface receptors may form a complex regulatory network that modulates T cell activity.

To further explore the physiological role of B7-H4 binding to Gal-9 *in vivo*, Gal-9 KO mice, B7-H4 KO mice, and B7-H4/Gal-9 DKO mice were generated (**Fig. S6A–E**) and were phenotyped for splenic immune cell proportions. Adult Gal-9 KO mice exhibited a significant increase in the proportion of CD4^+^ T cells and a significant decrease in the proportion of CD8^+^ T cells compared to WT mice, whereas no significant difference was observed in the proportions of total T cells (**Fig. 5A-B**), as well as B cells, NK cells, or CD11b^+^ cells between Gal-9 KO and WT mice (**Fig. S6F–G)**. These results indicate that CD4^+^ and CD8^+^ T cells are the primary immune lineages affected by Gal-9 deficiency in mice. Considering that Gal-9 induces T cell death *in vitro*, and the total T cell proportion remains unchanged, the observed reduction in the CD8^+^ T cell proportion is likely an indirect effect caused by the increased proportions of CD4^+^ T cells.

**Fig. 5.**
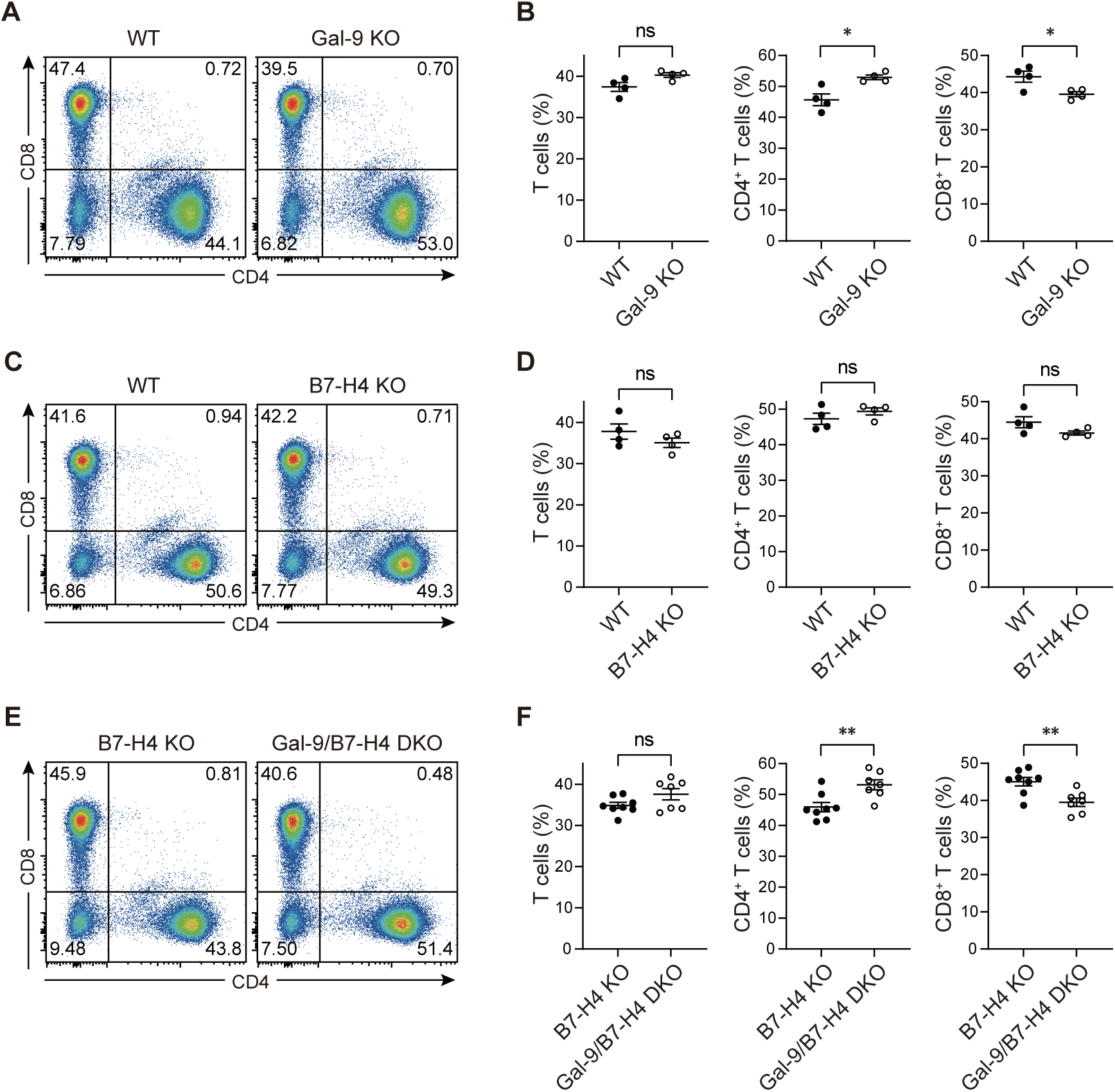
Gal-9 knockout increases the proportion of splenic CD4^+^ T cells in mice. (A-B) The proportion of splenic CD4^+^ T cells or CD8^+^ T cells in Gal-9 KO mice compared to those in WT mice by flow cytometry. Representative dot plots (gated on CD45^+^ live cells; see Fig. S6F) from one mouse are shown in panel A. The data from all mice, along with statistical analysis of splenic T cells, CD4^+^ T cells, or CD8^+^ T cells in Gal-9 KO mice compared to WT mice, are presented in panel B. n = 4 mice each group. (**C-D)** The proportions of splenic CD4^+^ and CD8^+^ T cells in B7-H4 KO mice compared to those in WT mice by flow cytometry. CD45^+^ live cell analysis is shown in Fig. S6G. Panels C and D are laid out similarly to Panels A and B, respectively. n = 4 mice each group. (**E-F)** Splenic T cell subsets in B7-H4/Gal-9 DKO mice compared to those in B7-H4 KO mice by flow cytometry. Panels E and F are laid out similarly to Panels A and B, respectively. n = 8 mice for B7-H4 KO group, n = 7 mice for B7-H4/Gal-9 DKO group. The unpaired t-test (panels B, D, F) is used for the two-group comparisons. All data are presented as mean ± s.e.m.

Notably, adult B7-H4 KO mice exhibited no significant changes in the proportions of the six immune cell lineages examined—total T cells, CD4^+^ T cells, CD8^+^ T cells, B cells, NK cells and CD11b^+^ cells—compared to WT mice (**Fig. 5C–D, Fig. S6H–I**). Further analysis of B7-H4/Gal-9 DKO mice revealed a significant increase in the proportion of CD4^+^ T cells and a significant decrease in the proportion of CD8^+^ T cells compared to B7-H4 KO mice (**Fig. 5E–F**), similar to what was observed in Gal-9 KO mice. These results suggest that B7-H4 KO alone is insufficient to alter Gal-9’s activity in regulating splenic immune cell composition *in vivo*. This is consistent with the earlier observation that multiple B7 family members and immune receptors on T cells interact with Gal-9.

Additionally, we examined the role of host tissue-expressed B7-H4 and Gal-9 in tumor progression by assessing tumor growth in B7-H4 KO, Gal-9 KO, and B7-H4/Gal-9 DKO mice. Using the murine colon adenocarcinoma cell line MC38, which lacks cell-surface mB7-H4 expression (**Fig. S7A**)—thereby precluding recognition of B7-H4 as a foreign antigen in B7-H4 KO or B7-H4/Gal-9 DKO mice—we found no difference in MC38 tumor growth among the three KO groups (**Fig. S7B**). These results indicate that host-expressed B7-H4 and Gal-9 are not critical for tumor progression in these models.

Intriguingly, contrary to previous reports suggesting tumor cell-expressed B7-H4 promotes tumor growth in WT immunocompetent mice (32), we found that mB7-H4-overexpressing MC38 cells (MC38-mB7-H4) (**Fig. S7A**) exhibited significantly reduced tumor growth in WT mice (**Fig. S7C**). This observation suggests that mB7-H4 overexpression on tumor cells does not promote tumor growth. Instead, it may activate the mouse immune response by interacting with unknown receptors or ligands, thereby enhancing antitumor activity. Furthermore, MC38-mB7-H4 cells failed to establish tumors in B7-H4 KO mice (**Fig.S7D**), suggesting mouse B7-H4 is recognized as a foreign antigen in the B7-H4 KO mice. Notably, MC38-mB7-H4 cells exhibited similar growth patterns to MC38 in two immunodeficient mouse models, with no significant differences observed in tumor growth curves (**Fig. S7E–F**). Additionally, by comparing the tumor growth curve in the immunodeficient mouse models (**Fig. S7E–F**) with those in the immunocompetent mouse model (**Fig. S7C**), it is further suggested that T cells in mice may play a critical role in mediating the antitumor activity triggered by mB7-H4 overexpression on tumor cells.

Collectively, our findings indicate that neither host-expressed B7-H4 nor tumor-expressed B7-H4 is essential for tumor progression in the contexts described above.

## Discussion

In this study, we identified Gal-9 as a previously unknown binding partner for B7-H4. Additionally, we uncovered several other previously unrecognized Gal-9 binding partners, including T cell surface receptors and multiple B7 family members. The molecular interaction between Gal-9 and B7-H4 is predominantly mediated by the glycan in the IgC domain of B7-H4 and the R65 residue in Gal-9’s N-CRD domain. Similarly, other binding partners also rely on their glycosylation for Gal-9 binding, while the R65 residue in Gal-9’s N-CRD domain is either dispensable or essential, depending on the binding partner. Moreover, both soluble and cell-surface B7-H4 counteract with Gal-9-induced T cell death and Gal-9-induced CD28 activation *in vitro*. While Gal-9 KO in mice resulted in an increase in CD4^+^ T cells within the splenic immune cell composition, B7-H4 KO had no effect on splenic immune cell composition. The DKO of B7-H4 and Gal-9 showed no additional effect compared to the *in vivo* activity observed with Gal-9 KO alone. Furthermore, Gal-9 KO, B7-H4 KO, or DKO of both Gal-9 and B7-H4 did not alter anti-tumor immunity in MC38 tumor-bearing C57BL/6 mouse model. Our findings indicate that it is likely B7-H4, in conjunction with other B7 family members and T cell surface immune receptors, interacts with Gal-9 as part of a complex regulatory network that finely modulates T cell activity through multiple and overlapping pathways.

The identification of B7-H4’s binding partner or functional receptor has been challenging due to the lack of B7-H4 binding target cells. It is worth noting that, in this study, we identified Gal-9 as a binding partner for B7-H4 using peritoneal immune cells isolated from mice adoptively transferred with OVA-expressing EG7 tumor cells and OVA-specific OT-1 T cells as the target cells bound by B7-H4 in the IP-MS method. It is apparent that Gal-9 is either induced to be expressed or markedly upregulated in peritoneal immune cells following the adoptive transfer of EG7 and OT-1 cells in mice, though the mechanism underlying this induction remains unknown. Notably, the peritoneal OT-1 cells themselves did not bind to B7-H4; instead, it was the host peritoneal immune cells bound to B7-H4. We surmise that the immune response, not necessarily limited to the OVA-specific OT-1 T cell response, following adoptive transferring of both tumor cells and OT-1 cells, induced upregulation of Gal-9 expression in peritoneal host immune cells. Indeed, previous reports indicate that Gal-9’s expression can be induced by various factors, such as IFNγ, IL-1β, and TNFα (20).

Interestingly, B7-H4, a member of the B7 family and the immunoglobulin superfamily, has been proposed to be a glycosylphosphatidylinositol (GPI)-anchored cell surface membrane protein (3, 33). Its protein expression is undetectable on immune cells in both humans and mice, even following stimulation, which is consistent with what we observed in this study. Additionally, B7-H4 exhibits low to undetectable protein expression levels in most healthy tissues. Notably, B7-H4 protein is preferentially overexpressed in tumor cells across various human cancer tissues. Such an expression pattern differs from other B7-CD28 family members. For instance, B7.1, B7.2, CTLA-4, PD-1, and PD-L1 are expressed on antigen-presenting cells or resting T cells, or they are induced after stimulation. In contrast, B7-H4 exhibits a more extensive expression profile in human cancers. Structurally, the human B7-H4 IgV domain adopts a typical IgV organization, as revealed by crystal structure studies (34), whereas the structure of its IgC domain remains unresolved. For B7-CD28 family members, the receptor-binding interface is generally located in the IgV domain. However, in this study, we found it is the IgC domain of B7-H4 that interacts with Gal-9, not the IgV domain.

Even though our findings does not rule out the possibility that the IgV domain of B7-H4 binds to an as-yet unidentified receptor involved in immune-regulating functions—or that it may potentially interacts with such a receptor through its IgV domain—our results, including the lack of an observable immune cell phenotype and the absence of an impact on anti-tumor responses against tumor challenge in B7-H4 deficiency in mice compared to WT mice, at least partially align with prior research (35, 36). These studies indicate that the impact of B7-H4 deficiency is minimal in mice on both C57BL/6 and BALB/c backgrounds. Similar results were observed in cancer models: B7-H4 KO in BALB/c mice challenged with mammary carcinoma cells (4T1) showed no differences in primary tumor growth (37). However, in the 4T1 metastatic model, B7-H4 KO BALB/c mice exhibited a reduced lung metastasis burden (36). Taken together, we surmise that B7-H4 likely plays a redundant and/or context-dependent role in immune regulation.

The context-dependent role for B7-H4 in mice has also been highlighted in other disease models observed in B7-H4 KO mice under conditions of additional challenges. In a Streptococcus pneumoniae infection model, B7-H4 deficient mice exhibited reduced disease severity, increased activation of CD4^+^ and CD8^+^ T cells in the lungs (38), suggesting a role for B7-H4 in suppressing T cell immunity. In autoimmune models, B7-H4 deficiency was associated with worsened disease outcomes (39). For example, severe experimental autoimmune encephalomyelitis was marked by increased Th1 and Th17 responses in pancreatic islets and the central nervous system. In a kidney disease model, B7-H4 deficiency led to severe renal injury, enhanced humoral responses, and polarization of inflammatory macrophages (40). These findings suggest that B7-H4 plays a regulatory role in immune responses across both infection and autoimmunity. However, whether these effects are mediated through an unidentified receptor interacting with its IgV domain or through Gal-9 binding to its IgC domain remains unclear.

Additionally, given our observation that Gal-9 interacts with multiple glycosylated protein partners, enabling context-dependent regulation, it is likely that B7-H4 functions as one of several negative co-signaling molecules that collectively fine-tune T cell-mediated immune responses within this context. Specifically, in this study, we found that several immune receptors, including CD28, 2B4, CD226, and SLAMF1, bind to Gal-9 at levels comparable to PD-1, TIM-3, and B7-H4. CD28 interacts with Gal-9 in a manner similar to its interaction with TIM-3, with the binding primarily mediated by the C-CRD domain of Gal-9, whereas the N-CRD of Gal-9 interacts with B7-H4. Other B7 family members, such as B7-H2, B7.1, B7-DC, and B7.2, were also found to bind to Gal-9 in our study. Previous studies also reported other Gal-9 binding partners, including TIM-3, PD-1, CD44, CD40, 4-1BB, death receptor 3 (DR3), V-domain Ig containing suppressor of T cell activation (VISTA), TLR-4, CD44, Dectin-1, Dectin-2, and ERMAP (41, 42). Given such a complex interaction network surrounding Gal-9, it is unsurprising that B7-H4 KO alone in mice does not result in immune cell phenotypes similar to those observed in Gal-9 KO mice.

Gal-9, a tandem-repeat type galectin consisting of two homologous but distinct CRDs (N-CRD and C-CRD), is expressed in almost all organs and plays a role in a variety of physiological functions, including cell growth, differentiation, adhesion, and cell death (43). Its cellular location is often intracellular but can also be extracellular at the cell surface or within the extracellular matrix (20, 44). Within the immune system, Gal-9 is widely expressed in immune cells, with accumulating evidence demonstrating it acts as a pleiotropic immune modulator. As mentioned above, Gal-9 interacts with multiple glycosylated immune cell surface receptors and B7 family members, including both previously known ones and the new ones identified in this study, highlighting the complexity of its role in immune regulation. The variable expression of these receptors across cell types likely influences Gal-9 signaling outcomes. Additionally, Gal-9’s ability to form lattices, such as TIM-3-Gal-9-PD-1 (19), adds another layer of possible regulation mechanism depending on the co-expression of other receptors.

In this study, since B7-H4 was identified as a binding partner of Gal-9, we further investigated B7-H4’s effect on Gal-9’s activity. Given the aforementioned complex and multifaceted nature of Gal-9’s activity, we specifically focused on its well-known dual immune-regulatory roles: stimulating or suppressing T cell activity, depending on its cellular location and the conditions (20–22). Intracellular Gal-9 has been shown to translocate to the membrane during the immune synapse formation upon stimulation, and Gal-9 deficiency in T cells impaired the phosphorylation of proximal TCR signaling proteins, suggesting that Gal-9 may participate in the modulation of TCR signals (21). Notably, we found that Gal-9 binds to CD28, a pivotal co-stimulatory receptor that enhances TCR signaling. Furthermore, we observed that Gal-9 binding to CD28 on T cells stimulated CD28 signaling activation in a manner similar to B7.1. Gal-9 also exhibited additive effects with B7.1 in enhancing this activation. Additionally, B7-H4, by binding to Gal-9, interferes with this signaling process and inhibits Gal-9-induced CD28 activation in T cells in an IgC domain and glycosylation dependent manner. Our findings provide at least one mechanism that may explain Gal-9-mediated T cell stimulation.

On the other hand, exogenous soluble Gal-9 is known for its role in inducing T cell apoptosis (16, 23, 26), suggesting that extracellular Gal-9 *in vivo* may also exert this effect, triggering apoptosis in T cells. Supporting this notion, we found that Gal-9 KO mice exhibited a significant increase in the proportion of splenic CD4^+^ T cells. *In vitro*, we also found that soluble Gal-9-induced T cell death in primary T cells and other T cell lines, and B7-H4 reduced this process by binding to Gal-9 in a glycosylation-dependent manner. Research has been conducted to elucidate the mechanisms underlying Gal-9-induced T cell death, with TIM-3- or VISTA-mediated mechanisms being reported (16, 45). The finding that B7-H4 inhibits Gal-9-induced T cell death appears contradictory to B7-H4’s known inhibitory immune-regulating role. However, this effect may resemble the process in which PD-1 binding to Gal-9, reducing TIM-3-dependent T cell death (19) and thereby contributing to the persistence of specific subset of exhausted T cells. Alternatively, B7-H4 may compete with TIM-3 or other Gal-9 binding partners on T cells to attenuate Gal-9-induced T cell death, reflecting a complex network of immune regulation that relies on the balanced interplay of activating and suppressing signals, as well as spatial and temporal dynamics.

In summary, our study identifies Gal-9 as a binding partner of B7-H4, along with a number of previously unknown Gal-9 binding partners. Gal-9 interacts with the co-stimulatory receptor CD28 on human T cells, leading to enhanced activation of downstream signaling. Additionally, Gal-9 induces T cell death *in vitro*, a finding supported by *in vivo* observations where Gal-9 deficiency in mice drives a significant increase in the proportion of splenic CD4^+^ T cells, underscoring the role of Gal-9 in T cell death. B7-H4 binding to Gal-9 reduces these Gal-9’s activities, offering novel insights into the molecular mechanism by which B7-H4 contributes to the T cell-mediated immune regulations. Nevertheless, the physiological and pathological relevance of these findings has yet to be fully elucidated and warrants further investigation.

## Methods

### Cell lines and primary immune cells

Jurkat, MOLT-4, 293T, 4T1, MC38 and EG7 cells were from the Cell Bank of Type Culture Collection (Chinese Academy of Sciences) or the American Type Culture Collection (ATCC). Freestyle 293F cells were from Thermo Fisher Scientific. MGAT1 (encoding GnT1) gene KO 293F cells were previously generated in our laboratory (46). Jurkat and MOLT-4 T cells were cultured in RPMI-1640 medium supplemented with 10% FBS. 293T cells were cultured in DMEM medium containing 10% FBS. 293F cells were cultured in SMM 293-TII medium. 4T1 cells were cultured in ATCC-formulated RPMI-1640 medium supplemented with 10% FBS. MC38 cells were cultured in DMEM supplemented with 10% FBS. EG7 cells were cultured in ATCC-formulated RPMI-1640 supplemented with 0.05 mM 2-mercaptoethanol, 10% FBS, and 0.4 mg/mL of G418. Primary human T cells were isolated from frozen human PBMCs (ORIBIOTECH, FPB003F-C) using the EasySep™ Human T Cell Enrichment Kit (Miltenyi Biotec) and maintained in RPMI-1640 without serum. Mouse splenic immune cells from Gal-9 KO mice were isolated using Red Blood Cell lysis buffer (0.15 M NH_4_Cl, 10 mM KHCO_3_, 0.1 mM Na_2_EDTA).

### Establishment of stable cell lines

Jurkat-NFAT-Luc stable reporter cell line was generated by transfecting Jurkat T cells with a plasmid carrying NFAT-RE-luciferase reporter encoding gene (luciferase gene under the control of an NFAT-response element) following the instruction of Amaxa Cell Line Nucleofector Kit V. The transfected cells were selected with 7.5 µg/mL antibiotics Blasticidin S (Invivogen, ant-bl-05) and subjected to limiting dilution cloning to obtain single stable cell clone. To assess NFAT activation, the selected stable clones were incubated with test agents, and NFAT signaling activity was quantified by measuring luminescence using the Bright-Glo™ Luciferase Assay System Kit (Promega, E2620).

MC38-mB7-H4 stable cells were established by transducing MC38 cells with lentiviruses expressing the full-length mB7-H4 protein. The lentiviruses were produced by transient transfection of 293T cells with the pLKO.1-VSV-G packaging plasmids. Lentivirus-transduced MC38 cells were selected with 5 µg/mL puromycin (Amresco, J593-25MG) and subsequently sorted using flow cytometry to isolate single-cell clones stably expressing cell surface mB7-H4. A candidate MC38-mB7-H4 stable cell clone, which exhibited a proliferation rate comparable to that of parental MC38 cells, was selected for subsequent tumor inoculation studies in mice.

### Antibodies and reagents

Antibodies and reagents used for flow cytometry analysis were primarily obtained from BioLegend unless otherwise indicated: FITC anti-mouse CD45 (clone 30-F11), FITC anti-mouse CD45.1 (clone A20), APC-Cy7 anti-mouse CD45.2 (BD Biosciences, clone 104), PE-Cy7 anti-mouse CD3 (clone 17A2), PerCP-Cy5.5 anti-mouse CD19 (clone 6D5), BV786 anti-mouse CD11b (BD Biosciences, clone M1/70), BV605 anti-mouse CD4 (clone GK1.5), BV421 anti-mouse NK1.1 (clone PK136), APC anti-mouse CD8 (clone 53-6.7), LIVE/DEAD Fixable Dead Cell Stain (Thermo Fisher Scientific, L34976), PerCP anti-human CD3 (clone UCHT1), BV605 anti-human CD4 (clone OKT4), APC anti-human CD8 (clone HIT8a), PE-Cy7 anti-human CD14 (clone HCD14), FITC anti-human CD19 (clone HIB19), BV421 anti-human CD56 (clone HCD56). Antibodies and reagents used for western blotting were primarily obtained from ABclonal unless otherwise indicated: Phospho-CD28 Rabbit mAb (Cell Signaling Technology, 16399), CD28 Rabbit mAb (A20346), Phospho-Akt Rabbit mAb (AP1208), Pan-Akt Rabbit mAb (A18675), HRP-conjugated β-Actin Rabbit mAb (AC028). Protease/Phosphatase Inhibitor Cocktail (Cell Signaling Technology, 5872). Human Galectin-9 (Gal-9) protein was purchased from R&D Systems (Cat. 2045-GA). Other reagents used in this study are listed in **Table S1**.

### Protein expression, purification, and de-glycosylated protein preparation

The mB7-H4-Fc protein was expressed using an expression plasmid constructed with the cDNA encoding the mB7-H4 ectodomains, comprising the IgV and IgC domains (UniProt Q7TSP5, residues 32-261), fused to the human IgG1 Fc region. Similarly, expression plasmids for hB7-H4-Fc, hB7-H4 IgV-Fc, and hB7-H4 IgC-Fc proteins were constructed using the cDNAs encoding the hB7-H4 ectodomain (UniProt Q7Z7D3-1, residues 32-259), the IgV domain (UniProt Q7Z7D3-1, residues 36-148), and the IgC domain (UniProt Q7Z7D3-1, residues 153-259) respectively, fused to the mouse IgG2a Fc region. 293F cells were transiently transfected with these expression plasmids using PEI-MAX transfection reagents. Five days post-transfection, the culture supernatant was harvested and purified using Protein A beads. Protein concentrations were determined using a Nanodrop spectrophotometer. The de-glycosylated hB7-H4-Fc protein (B7-H4^MGAT1-^) was produced by transient transfection of MGAT1 KO 293F cells with the hB7-H4-Fc plasmid, and purified using Protein A affinity chromatography, similar to the hB7-H4-Fc described above.

The mB7-H4-Bio protein was expressed by co-transfecting 293F cells using PEI-MAX with two plasmids: an expression plasmid encoding mB7-H4 ECD-His_6_-Avi, the mouse B7-H4 extracellular domain (ECD; UniProt Q7TSP5, residues 1-261) fused to a C-terminal His_6_-Avi tag (GLNDIFEAQKIEWHE), and a BirA biotin ligase-expressing plasmid. Five days post-transfection, the culture supernatant was harvested and purified by Ni-NTA Agarose affinity chromatography (QIAGEN, 30230). Similarly, human Sema3a (UniProt Q14563, residues 1-771) with a C-terminal His_6_ tag was expressed in 293F cells and purified.

The recombinant proteins (Fc tag, mouse IgG2a isotype) of multiple human immune receptors, including 2B4 (UniProt Q9BZW8-2, residues 1-224), PD-1 (UniProt Q15116, residues 1-170), TIM-3 (UniProt Q8TDQ0-1, residues 1-202), CD226 (UniProt Q15762, residues 1-254), SLAMF1 (UniProt Q13291-1, residues 1-237), CD28 (UniProt P10747-1, residues 1-152), BTLA (UniProt Q7Z6A9-1, residues1-156), ICOS (UniProt Q9Y6W8-1, residues 1-140), TIGIT (UniProt Q495A1-1, residues 1-141), CD305 (UniProt Q6GTX8-1, residues 1-165), CD160 (UniProt O95971-1, residues 1-159), CLTA-4 (UniProt P16410-1, residues 1-161), CD2 (UniProt P06729, residues 1-209), B7.1 (UniProt P33681-1, residues 1-242), B7.2 (UniProt P42081-1, residues 1-247), B7-DC (UniProt Q9BQ51-1, residues 1-220), B7-H1 (UniProt Q9NZQ7-1, residues 1-238), B7-H2 (UniProt O75144-1, residues 1-256), and B7-H3 (UniProt Q5ZPR3-2, residues 1-248) were expressed and purified, similar to the hB7-H4-Fc described above.

To generate site-directed mutants of human B7-H4, PCR primers targeting specific mutation sites were designed using TAKARA Primer design tools and employed to amplify DNA fragments in accordance with the manual of High-fidelity DNA polymerase KOD-Plus (TOYOBO). Molecular cloning was carried out according to the manual of In-Fusion HD Cloning Kit (Takara). These B7-H4 mutants (Fc tag, mouse IgG2a isotype) were expressed and purified as described above.

For removing the N-linked glycosylation modifications and preparing of de-glycosylated hB7-H4-Fc, 10 µg of hB7-H4-Fc and 2 µL of PNGase F (New England Biolabs, P0704S) were mixed and then incubated for 24 h at 37 °C under non-denaturing reaction conditions. An untreated control (10 μg hB7-H4-Fc without PNGase F) was processed in parallel. To prepare the denaturing reaction using PNGase F, hB7-H4-Fc was first pre-denatured at 100 °C for 10 min. A mixture of 10 µg of pre-denatured protein, 2 µL of GlycoBuffer 2 (New England Biolabs), 2 µL of 10% NP-40, 6 µL of H_2_O, and 1 µL of PNGase F, was then incubated for 1 h at 37 °C. The treated protein samples were then subjected to SDS-PAGE electrophoresis and Coomassie blue staining for confirming the de-glycosylation.

### Jurkat-NFAT reporter bioassay

Jurkat-NFAT-Luc reporter stable cells were mixed with hB7-H4-Fc or Gal-9 proteins at indicated concentrations or mixed with 293F-B7-H4 cells (293F cells transiently expressing full-length hB7-H4), and incubated for 3 h at 37 °C in a 96-well plate. Cells were centrifuged (400 g, 5 min), resuspended in 70 μL of Glo Lysis Buffer (Promega, E2661), and incubated for 5 min. 50 µL of the cell lysate was transferred into a 96-well white bottom microplate and mixed with 50 µL of Bright-Glo Luciferase Assay Substrate (Promega, E2620). NFAT-mediated luminescence was measured using GloMax Navigator System (Promega).

### Surface Plasmon Resonance (SPR)

The binding kinetics between hB7-H4-Fc protein, its variants, and other Fc-tagged proteins to human Gal-9 were analyzed using SPR on a Biacore T200 instrument (Biacore, GE Healthcare). Protein A/G was immobilized on a CM5 sensor chip using an amine-coupling kit (GE Healthcare). hB7-H4-Fc or its variant proteins (1 μg/mL) were captured on the sensor chip, followed by the flow of two-fold serially diluted Gal-9 starting at 33 nM. Binding kinetics were analyzed using Biacore T200 evaluation software to calculate the association rate constant (ka), dissociation rate constant (kd), and equilibrium dissociation constant (KD).

### Immunoprecipitation-mass spectrometry (IP-MS)

For immunoprecipitation (IP), 50 µL of M-280 streptavidin magnetic beads were incubated with 1 mL cell lysate supernatant supplemented with or without 4 µg mB7-H4-Bio bait protein overnight at 4 °C. Following incubation, the beads were washed four times with PBST buffer (PBS containing 0.05 % Tween-20). The washed M-280 magnetic beads carrying the precipitated proteins were subjected to sodium dodecyl sulfate–polyacrylamide gel electrophoresis (SDS-PAGE). Silver staining analyses were conducted to identify candidate binding proteins for mB7-H4. Protein bands of interest, which were observed in the experimental lane with mB7-H4-Bio but absent in the control lane (without the bait protein), were excised and subjected to LC-MS/MS analysis at the Proteomics Facility of National institute of Biological Sciences, Beijing (NIBS).

### Western blotting

For immunoprecipitated (IP) samples, the precipitated proteins were resolved by SDS-PAGE and transferred to a nitrocellulose (NC) membrane using a wet transfer method. The NC membrane was blocked with 3% (w/v) non-fat milk in PBST for 1 h at room temperature, followed by incubation with HRP-Streptavidin (Thermo Fisher Scientific, 21130) in blocking buffer for 1 h at 37 °C. After five washes with PBST, the membrane was immersed into ECL substrate for 5 min. All blots on the x-ray films were processed inside the darkroom.

For cell lysates samples, the whole-cell lysates were mixed with SDS buffer at room temperature for 30 min, heated at 100 °C for 10 min, and centrifuged at 12,000 g for 10 min. Supernatants (4-8 µL) were separated by SDS-PAGE and transferred onto NC membranes. Membranes were incubated overnight at 4 °C using these primary antibodies, including Phospho-CD28 Rabbit mAb (Cell Signaling Technology, 16399), CD28 Rabbit mAb (ABclonal, A20346), Phospho-Akt Rabbit mAb (AP1208), or Pan-Akt Rabbit mAb (A18675). Subsequent blotting was processed as described above.

### Gal-9-mediated CD28/AKT activation

To examine Gal-9-mediated stimulation of pCD28 signaling activity in the presence of plate-bound B7.1, serum-starved Jurkat T cells (1.0 × 10^5^ cells per well) were treated with Gal-9 (3-7 µg/mL) in a 96-well plate pre-coated with 10 µg/mL hB7.1-Fc for 2 h at 37 °C. Following the incubation for 30 min at 37 °C, the plate was centrifuged at 400 g for 5 min to harvest cell pellets. The resulting cell pellets were lysed in RIPA buffer containing protease and phosphatase inhibitors for 30 min at 4 °C. The cell lysates were analyzed by western blotting as mentioned above. To assess the regulation of Gal-9-mediated stimulation of CD28 signaling by glycosylated B7-H4, serum-starved T cells (Jurkat, MOLT-4, or primary T cells; 1.0 × 10⁵ cells per well) were treated with 8 µg/mL Gal-9 for 30 min at 37 °C in the presence or absence of 25 µg/mL glycosylated hB7-H4-Fc, hB7-H4 IgV-Fc, or de-glycosylated B7-H4^MGAT1-^ fusion proteins. After centrifugation at 400 g for 5 min to cell pellets, the cell lysates were prepared and analyzed by western blotting as described above.

### Cell viability assay

Serum-starved MOLT-4, Jurkat, or primary T cells (1.0 × 10^5^ cells/sample) were filtered through 40 µm strainers and treated with 8 µg/mL Gal-9 in the presence or absence of 25 µg/mL glycosylated hB7-H4-Fc or de-glycosylated B7-H4^MGAT1-^ for 30 min. Following the treatment, 7-AAD was added to the T cells and incubated for 5 min. Flow cytometry data were acquired using the BD Accuri C6 instrument and analyzed using Flowjo (v10.6.2). The analysis focused on determining the proportion of live T cells (7-AAD negative) within the parent gate.

To examine the regulation of T cell viability by immobilized B7-H4 proteins, glycosylated B7-H4 or deglycosylated B7-H4^MGAT1-^ (8 µg/mL) were immobilized overnight on Protein A beads (Smart-Lifesciences, SA023005). The next day, 1.0 × 10^5^ of serum-starved MOLT-4 T cells were incubated for 30 min at 37 °C with Gal-9 (6 µg/mL) and either iB7-H4-or iB7-H4^MGAT1-^-immobilized beads. Cells were then stained with 7-AAD for 5 min and analyzed by flow cytometry as previously described.

To assess Gal-9 effects on PBMC viability, human PBMCs (5 × 10⁵/well) were seeded in a 96-well plate and treated with or without 8 µg/mL Gal-9 for 30 min. After washing twice with PBS, cells were stained with LIVE/DEAD^TM^ Fixable Near-IR dye (Thermo Fisher Scientific, L34975) for 30 min at 4 °C. Following this, the cells were washed twice with 0.5% BSA/PBS, and incubated with primary antibodies targeting human immune lineages for 15 min at 4 °C. Following two additional washes, cells were resuspended in 300 µL of PBS for flow cytometry data acquisition on a BD LSRFortessa™. Data were analyzed using FlowJo (v10.6.2). Viability of CD3^+^ T cells and CD19^+^ B cells within live PBMCs were quantified using GraphPad Prism (v9.2.0).

### Flow cytometry

For examining the binding of mB7-H4 to host CD45.1^+^ immune cells or CD45.1^-^ OT-1 T cells harvested from the peritoneal cavity of adoptive-transfer-model mice, the harvested immune cells were stained with an amino-reactive dead cell dye (Thermo Fisher Scientific, L34975) for 30 min at 4 °C. After two washes with 0.5% BSA/PBS, cells were incubated with or without mB7-H4-Fc for 30 min at 4 °C. Following two additional washes, cells were stained with: PE anti-human Fc antibody (Thermo Fisher Scientific, 12-4998-82), APC-Cy7 anti-mouse CD45.2 (BD Biosciences, clone 104), FITC anti-mouse CD45 (BioLegend, clone 30-F11), or FITC anti-mouse CD45.1 (BioLegend, clone A20) for 15 min at 4 °C. Following another two washes, these cells were resuspended in 300 µL of PBS for flow cytometry data acquisition on a BD LSRFortessa^TM^. All flow cytometry data were analyzed with Flowjo (v10.6.2).

To analyze immune cell subsets in spleens of Gal-9 KO mice, the spleens were processed into single-cell suspensions. After centrifugation at 300 g for 5 min, the splenic cells from half of the suspension were treated with Red Blood Cell lysis buffer to lyse red blood cells. Following this, the splenocytes were then filtered through a 40 µm cell strainer, and stained with an amino-reactive dead cell dye (Thermo Fisher Scientific, L34975) for 30 min at 4 °C. Following two washes with 0.5% BSA/PBS, cells were incubated with the primary antibodies targeting mouse immune lineages for 15 min at 4 °C. The flow cytometry data were acquired and analyzed as described above. Proportions of immune subsets within each parent gate were statistically analyzed using GraphPad Prism (v9.2.0).

The interaction between Fc-tagged proteins (hB7-H4-Fc variants/others) and cell-expressed Gal-9 or its variants were analyzed using transiently transfected 293T cells. The 293T cells were transfected to express either the full-length Gal-9 (UniProt O00182-1, residues 1-355, including both the N-CRD and C-CRD domains) or its truncation/mutation variants, each fused with EGFP. The transfected 293T cells were fixed and permeabilized following the instructions of the Fixation/Permeabilization Kit (BD). Subsequently, these cells were incubated with 10 µg/mL test proteins for 30 min at 4 °C. After washing twice with Perm/Wash Buffer (BD), the cells were stained with Alexa Fluor 647 anti-mouse Fc antibody (Thermo Fisher Scientific, 1839633) for 20 min at 4 °C. Following two additional washes, the cells were resuspended in PBS and analyzed by flow cytometry. Flow cytometry data were acquired and analyzed as described above. The interaction of cell-expressed Gal-9 with multiple human immune receptors (hPD-1-Fc, hTIM-3-Fc, hCD28-Fc, h2B4-Fc, hSLAMF1-Fc, hBTLA-Fc, hICOS-Fc, hTIGIT-Fc, hCD305-Fc, hCD160-Fc, hCTLA-4-Fc, hCD2-Fc) or additional B7 family members (hB7.1-Fc, hB7.2-Fc, hB7-DC-Fc, hB7-H1-Fc, hB7-H2-Fc, hB7-H3-Fc) was similarly measured following the aforementioned protocols for hB7-H4-Fc.

For analyzing B7-H4-EGFP expression in transiently transfected 293F cells, a plasmid encoding full-length hB7-H4 (UniProt Q7Z7D3-1, residues 1-282), which includes the extracellular, transmembrane, and cytoplasmic domains, fused to an EGFP tag was constructed. 293F cells were transiently transfected with the plasmid using PEI-MAX transfection reagent. After two days of post-transfection, 5 × 10^5^ WT 293F and B7-H4-EGFP-expressing 293F cells were resuspended with 100 µL of PBS. Expression was quantified by flow cytometry as previously described.

For analyzing the binding of B7-H4 to activate human T cells, primary human T cells were activated with anti-human CD3/CD28 beads (Absin, abs160019) and cultured in RPMI-1640 medium containing 10% FBS and 1000 IU/mL IL-2. The activated human T cells were first stained with an amino-reactive dead cell dye (Thermo Fisher Scientific, L34975), and then incubated with hB7-H4-Fc or isotype control (human CD147-Fc) for 30 min at 4 °C. Following two washes with 0.5% BSA/PBS, the cells were stained with Alexa Fluor 647 anti-mouse Fc antibody for 20 min at 4 °C. Flow cytometry data acquisition and analysis were performed as described earlier. The binding of mouse B7-H4 to naive OT-1 T cells harvested from OT-1 mouse splenocytes and OVA-activated OT-1 T cells (cultured with 2 µg/mL OVA for 3 days) was similarly conducted by flow cytometry analysis as mentioned above.

### Animal study

C57BL/6J mice (CD45.1^+^, CD45.2^-^) were obtained from Jackson Laboratory. BALB/c Nude mice, CB-17 SCID mice, and wild-type (WT) mice (C57BL/6J background) were obtained from Charles River. OT-1 mice were provided by Dr. Yulu Li at NIBS. Gal-9 KO mice, B7-H4 KO mice, and B7-H4/Gal-9 DKO mice were generated on a C57BL/6J background at the Transgenic Animal Center of NIBS. All mice were maintained under specific-pathogen-free conditions in the Animal Facility of NIBS. The animal experiments were conducted in accordance with the approved protocols of the Institutional Animal Care and Use Committee of NIBS (NIBS2022M0038).

For establishment of the adoptive transfer mouse model, naive OT-1 T cells (CD45.1^-^, CD45.2^+^) were isolated from OT-1 mouse splenocytes using red blood cell lysis buffer. Cells were activated with 2 µg/mL OVA peptide for 3 days. Then OVA-activated OT-1 T cells (6.7 × 10^6^) were injected into the peritoneal cavity of C57BL/6J mice (CD45.1^+^, CD45.2^-^). On the next day (day 0), 7 × 10^7^ EG7 tumor cells isolated from EG7 tumor-bearing C57BL/6J mice using Tumor Dissociation Kit (Miltenyi, 130-096-730), were injected into the peritoneal cavity of C57BL/6J mice (CD45.1^+^, CD45.2^-^). On day 1, host CD45.1^+^ cells, OT-1 T cells, or EG7 tumor cells within the peritoneal cavity of these adoptive-transfer-model mice were harvested. The peritoneal immune cells were lysed with 1 mL of RIPA buffer (containing 1 mM proteinase PMSF) per up to 5 × 10^6^ cells for 45 min at 4 °C. Lysates were centrifugated at 12,000 g for 10 min. Aliquots of the supernatant were flash-frozen in liquid nitrogen and stored at -80 °C for use in the next step of the IP procedure.

Gal-9 KO mice were obtained using cryopreserved sperm from Gal-9 KO mice (GemPharmatech) and *in vitro* fertilization was performed by Transgenic Animal Center at NIBS. Offspring were genotyped to confirm the Gal-9 KO genotype. The primers used for genotype PCR were as follows: F1 5’- TAGACTCCTACGTCCTGAGCATCCT-3’, R1 5’-ATCCAGATCAGGCAGCTCCTAAC-3’, F2 5’- CTTGTGTTTGCTTGCTTCATGC-3’, R2 5’-CTAGGACTTGCTTGTTAGGCAAGC-3’.

For generation of B7-H4 KO mice, Exon3 and Exon4, which encode the mouse B7-H4 IgV and IgC ectodomains, were selected as the KO region. The gRNAs were designed and synthesized by GENEWIZ. Microinjection of the gRNAs and CRISPR/Cas9 vector into mouse fertilized eggs was performed by Transgenic Animal Center at NIBS. Genotype primers were designed based on the different genomic sequences between WT mice and B7-H4 KO mice. Genotype PCR was conducted according to the manual of High-fidelity DNA polymerase KOD-Plus (TOYOBO) to confirm the B7-H4 KO genotype. The primers used for genotype PCR were as follows: F3 5’-GAGTTCCTTCATCATTCCAAGAAAGACAAAG-3’, R3 5’-TTGGTACCTACAGCAGATCTGTGCAC-3’, F4 5’-GAGTTCCTTCATCATTCCAAGAAAGACAAAG-3’, R4 5’- GGTATCTGATATGTAACCCCTGGAAAGAATTG-3’.

For generation of B7-H4/Gal-9 DKO mice, B7-H4 KO female mice were served as the strain background. Cryopreserved sperm from Gal-9 KO male mice was used for *in vitro* fertilization. The resulting offspring were genotyped to confirm the B7-H4/Gal-9 DKO genotype using the genotype primers previously described for B7- H4 KO mice and Gal-9 KO mice.

For syngeneic mouse tumor models, including MC38 and MC38-mB7-H4 tumor models, male mice aged 10-12 weeks (BALB/c Nude, CB-17 SCID, C57BL/6J WT, B7-H4 KO, Gal-9 KO, or B7-H4/Gal-9 DKO) were inoculated subcutaneously with 5 × 10^5^ tumor cells in the right lower flank (day 0). Tumor dimensions were measured every 4 days using calipers. Tumor volume was determined according to the following formula based on caliper measurements: tumor volume = 0.5 × length × width^2^. In accordance with Ethical Approval for Research Involving Animals at NIBS, the experimental observation was terminated when mouse tumor volume approached 1500 mm^3^.

### Statistical analysis

Statistical comparisons among three or more groups were determined using one-way ANOVA and Tukey’s multiple comparisons test, while two-group comparisons were determined by unpaired Student’s t-test. All analyses were performed using GraphPad Prism (v9.2.0). Significance levels were defined as follows: *P* > 0.05 indicates no significant (ns) effect on the group means, and these *P* values signifies that the mean of one group is significantly different from another, including * *P* < 0.05, ** *P* < 0.01, *** *P* < 0.01, **** *P* < 0.0001.

## Acknowledgments

We thank Jing Wang and Yong Luo at Antibody center of NIBS for their technical assistance during the initial phase of this project. We thank the Transgenic Animal Center at NIBS for generating knockout mice and the Proteomics Facility at NIBS for mass spectrometry analysis.

## Author contributions

R.Z.W. and J.S. conceptualized this study. R.Z.W. performed experiments and prepared the figures. All authors interpreted the results and drafted the manuscript. J.S. supervised the study.

## Supplemental Figure Legends

**Fig. S1. The identification of B7-H4 binding proteins. (A-C)** Flow cytometry analysis of B7-H4 binding to naïve and activated T cells. hB7-H4-Fc binding to activated human T cells (A); mB7-H4-Fc binding to naïve or cell-cultured OVA-activated OT-1 T cells (B); mB7-H4-Fc binding to OT-1 T cells (CD45.1^-^, CD45.2^+^) isolated from adoptive-transfer-model mice as described in Methods (C). Histogram overlays show the isotype control group (shaded gray), and potential binding signals stained with hB7-H4-Fc or mB7-H4-Fc (black line). These Fc tag proteins were tested at 10 μg/mL. **(D)** Immunoprecipitation using mB7-H4-Bio. Cell lysates from 4T1 tumor cells isolated from 4T1 tumor-bearing mice, EG7 tumor cells isolated from EG7 tumor-bearing mice, or CD45.1^+^ peritoneal immune cells isolated from the adoptive-transfer-model mice (as described in Methods and Fig. 1A) were precipitated using mB7-H4-Bio. The arrow indicates the ∼35 kDa band precipitated by mB7-H4-Bio in CD45.1^+^ peritoneal immune cell lysates but not in 4T1 or EG7 tumor cell lysates. **(E)** The protein sequence of mouse Gal-9 (UniProt O08573-2, residues 1-322, 36.5 kDa). The seven peptide sequences in red represent those identified by LC-MS/MS-based proteomic analysis of the 35 kDa protein band.

**Fig. S2. Characterization of the binding between de-glycosylated B7-H4 mutants and Gal-9. (A)** A schematic of the human B7-H4 ectodomains and their glycosylation sites. Seven glycosylation sites are marked by asterisks, and the sites required for the interaction with Gal-9 are highlighted in red. **(B)** SDS-PAGE analysis of glycosylated hB7-H4-Fc and its de-glycosylated forms or single-site mutants. **(C)** SPR analysis of Gal-9 binding to B7-H4 glycosylation mutants. To test its binding, 1.2 µg/mL Gal-9 was flowed through a Protein A/G-coated chip immobilized with individual B7-H4 mutants.

**Fig. S3. Gal-9 decreases T cell viability. (A)** Gal-9 reduces the cell viability of human PBMCs. Flow cytometry dot plots show the proportion of live human PBMCs after treatment with 8 µg/mL Gal-9 for 0.5 h. **(B)** Gal-9 reduces the cell viability of CD3^+^ T cells in PBMCs. Flow cytometry dot plots show CD3^+^ T cell or CD19^+^ B cell proportions within the parent gate of live PBMCs in Panel A. **(C-D)** Gal-9 reduces the cell viability of primary human T cells. Flow cytometry dot plots show the proportions of live cell (7-AAD negative) within serum-starved primary T cells after treatment with 8 µg/mL Gal-9 for 0.5 h (C) or 18 h (D). Statistical significance was determined by unpaired t-tests (Panels A-D), to assess live T cells treated with Gal-9 compared to those untreated. For each panel (A-D), data from three independent replicates (mean ± s.e.m.) are shown on the right.

**Fig. S4. Gal-9 interacts with multiple immune checkpoint receptors. (A)** SPR binding kinetics between Gal-9 and the ectodomains of ten immune checkpoint receptors. The SPR analysis used the same method as mentioned in Fig. 4A. **(B)** Flow cytometry analysis of the binding activities between human 2B4, CD226, and SLAMF1 and 293T cells transiently overexpressing either WT Gal-9 or single-site mutant Gal-9 R65A. The analysis used the same method as Fig. 4B.

**Fig. S5. Gal-9 interacts with B7 family members. (A)** SPR binding kinetics between Gal-9 and the ectodomains of seven members in the B7 family. The SPR analysis used the same method as Fig. 4A. **(B)** Flow cytometry analysis of the binding between the B7 family members and 293T cells transiently overexpressing WT Gal-9. Flow cytometry analysis used the same method as mentioned in Fig. 4B.

**Fig. S6. Genotypic and phenotypic characterization of splenic immune cells in Gal-9 KO mice and B7-H4 KO mice. (A-B)** Schematic diagram of Gal-9 KO (A) and B7-H4 KO (B) design in WT C57BL/6J mice. For Gal-9 KO, genotype primers, F1 and R1, are designed to identify WT allele containing intact mouse Gal-9 gene—the Exon2 (89 bp) of mouse Gal-9 gene, while F2 and R2 are designed to identify Gal-9 KO allele. For B7-H4 KO, genotype primers, F3 and R3, are designed to identify WT allele containing intact mouse B7-H4 gene, while F4 and R4 are designed to identify B7-H4 KO allele lacking the whole KO fragment (including Exon3, encoding for mouse B7-H4 IgV domain; and Exon4, encoding for mouse B7-H4 IgC domain). **(C-D)** Genotyping results of Gal-9 KO (C) and B7-H4 KO (D) newborn mice. The PCR products, ① and ②, correspond to the target genes amplified using the genotype primers as described above in Panel A; the PCR products, ③ and ④, correspond to the target genes amplified using the genotype primers as described above in Panel B. **(E)** Genotyping of B7-H4 KO and B7-H4/Gal-9 DKO newborn mice. The PCR products correspond to the target genes using all four pairs of genotype primers as described in Panels A-B. **(F)** Flow cytometry analysis of CD45^+^ live or dead splenic cells in Gal-9 KO mice compared to those in WT mice. These data are from the same experiment as described in Fig. 5A-B. Representative dot plots from one mouse are shown on the left. The data from all mice (presented as mean ± s.e.m.), along with the statistical analysis of splenic CD45^+^ live cells in Gal-9 KO mice compared to WT mice, are presented on the right (n = 4 mice for each group). **(G)** The data from all mice are shown in Panel F, along with the statistical analysis of splenic B cells, CD11b^+^ cells or NK cells in Gal-9 KO mice compared to WT mice. **(H-I)** These datasets as presented in Panels F-G were used to assess the splenic immune cells in B7-H4 KO mice compared to those in WT mice. The unpaired t-test was used for two-group statistical comparisons as described in Panels F-G.

**Fig. S7. MC38 tumor growth exhibits no difference between B7-H4 KO mice and Gal-9 KO mice. (A)** Flow cytometry analysis of B7-H4 expression on MC38-mB7-H4 stable cells. Histogram overlays show the isotype control antibody (shaded gray) and mB7-H4 expression detected using anti-mB7-H4 antibody (black line). **(B)** MC38 tumor growth in WT, Gal-9 KO, and B7-H4/Gal-9 DKO mice. The data shown on the left is from one experiment (n = 3 mice per group), while the data shown on the right is from another independent experiment, n = 5 mice per group, except for the WT C57BL/6J mice (n = 4). (**C-F)** Tumor growth comparison of MC38 and MC38-mB7-H4 in WT C57BL/6J mice (C), B7-H4 KO mice (D), CB-17 SCID (E), and BABL/c Nude mice (F). In each panel, MC38 and MC38-mB7-H4 tumor cells were implanted bilaterally in the same mouse (MC38 on the right flank and MC38-mB7-H4 on the left flank), with two groups sharing the same set of mice (n = 4 mice per group). The two-way ANOVA tests were used to determine the statistical significance between groups. All data are presented as means ± s.e.m.

