## supplemental figure and table for "B7-H4 Binds Galectin-9 Glycosylation-Dependently and Attenuates Galectin-9-Mediated CD28/AKT Activation and T Cell Death"

### Fig. S1

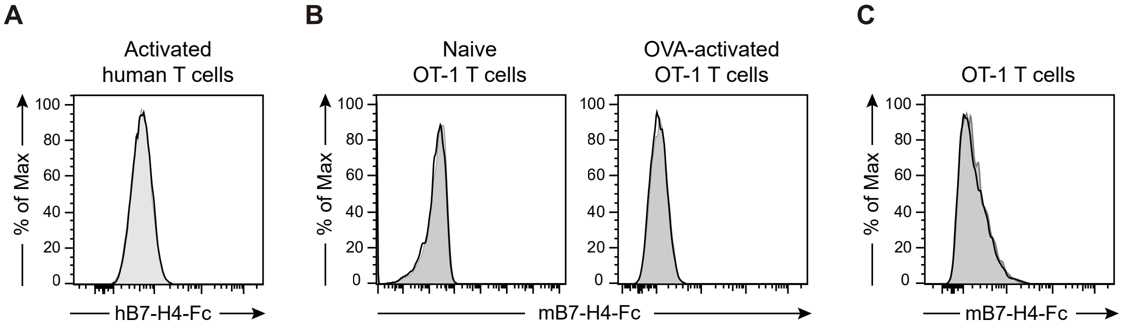


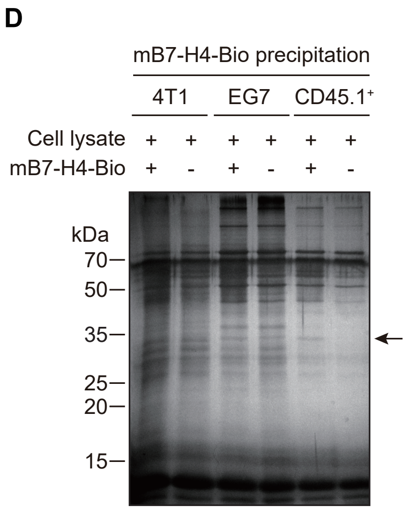


**E**

Protein sequence of mouse Galectin-9 (36.5 kDa)

| | | | |
|---|---|---|---|
| 1 | MALFSAQSPY | INPIIPFTGP | IQGGLQEGLQ |
| 31 | VTLQGTTKSF | AQRFVVNFQN | SFNGNDIAFH |
| 61 | FNPRFEEGGY | VVCNTKQNGQ | WGPEERKMQM |
| 91 | PFQKGMPFEL | CFLVQRSEFK | VMVNKKFFVQ |
| 121 | YQHRVPYHLV | DTIAVSGCLK | LSFITFQTQN |
| 151 | FRPAHQAPMA | QTTIHMVHS | TPGQMFSTPGI |
| 181 | PPVVYPTPAY | TIPFYTPIPN | GLYPSKSIMI |
| 211 | SGNVLPDATR | FHINLRCGGD | IAFHLNPRFN |
| 241 | ENAVVRNTQI | NNSWGQEERS | LLGRMPFSRG |
| 271 | QSFSVWIICE | GHCFKVAVNG | QHMCEYYHRL |
| 301 | KNLQDINTLE | VAGDIQLTHV | QT |

### Fig. S2

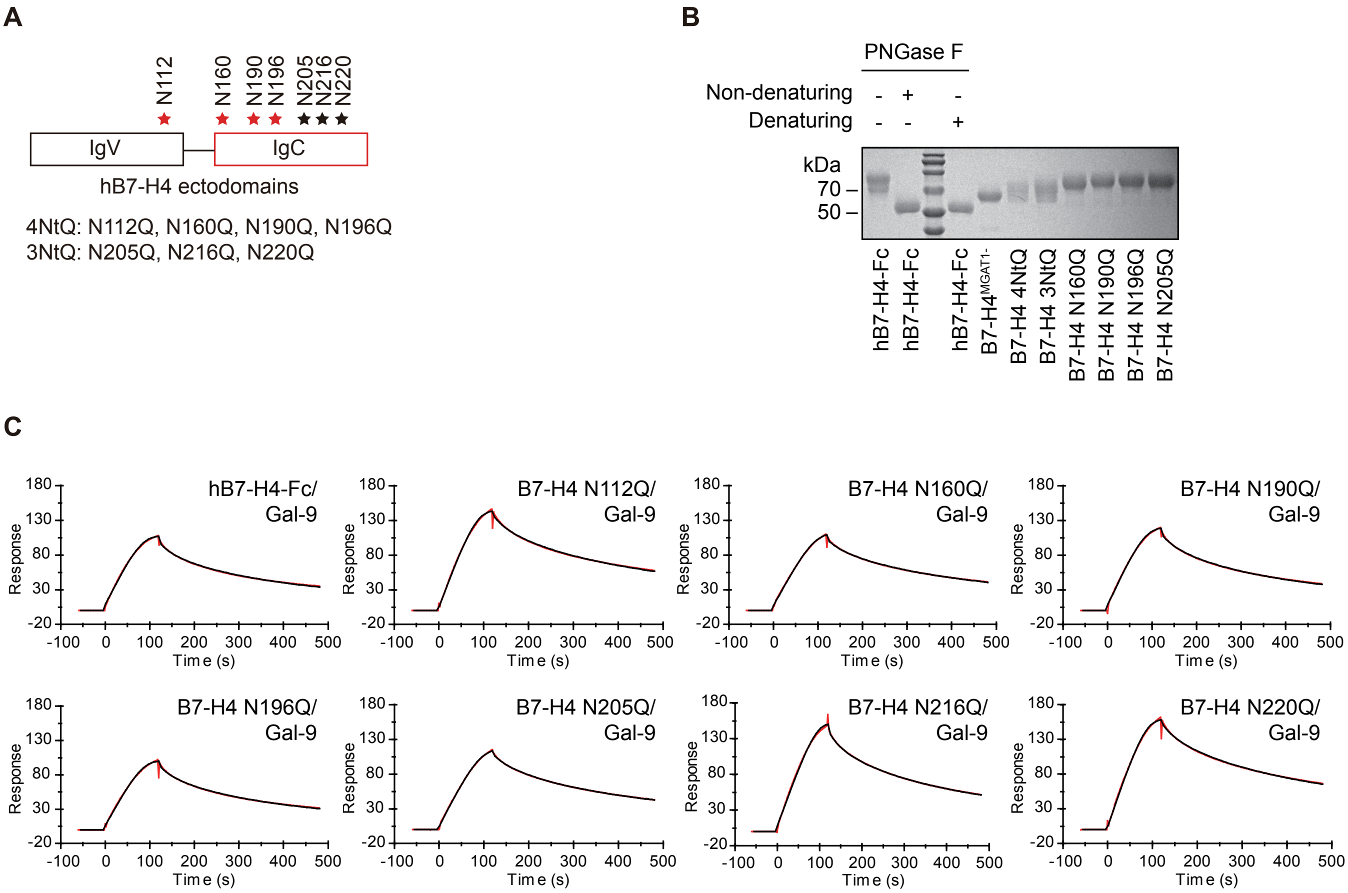
A
N112
N160
N190
N196
N205
N216
N220
IgV
IgC
hB7-H4 ectodomains
4NtQ: N112Q, N160Q, N190Q, N196Q
3NtQ: N205Q, N216Q, N220Q
B
PNGase F
Non-denaturing - + -
Denaturing - - +
kDa
70
50
hB7-H4-Fc
hB7-H4-Fc
hB7-H4-Fc
B7-H4MGAT1-
B7-H4 4NtQ
B7-H4 3NtQ
B7-H4 N160Q
B7-H4 N190Q
B7-H4 N196Q
B7-H4 N205Q
C
hB7-H4-Fc/Gal-9
B7-H4 N112Q/Gal-9
B7-H4 N160Q/Gal-9
B7-H4 N190Q/Gal-9
B7-H4 N196Q/Gal-9
B7-H4 N205Q/Gal-9
B7-H4 N216Q/Gal-9
B7-H4 N220Q/Gal-9
Response
Time (s)

### Fig. S3

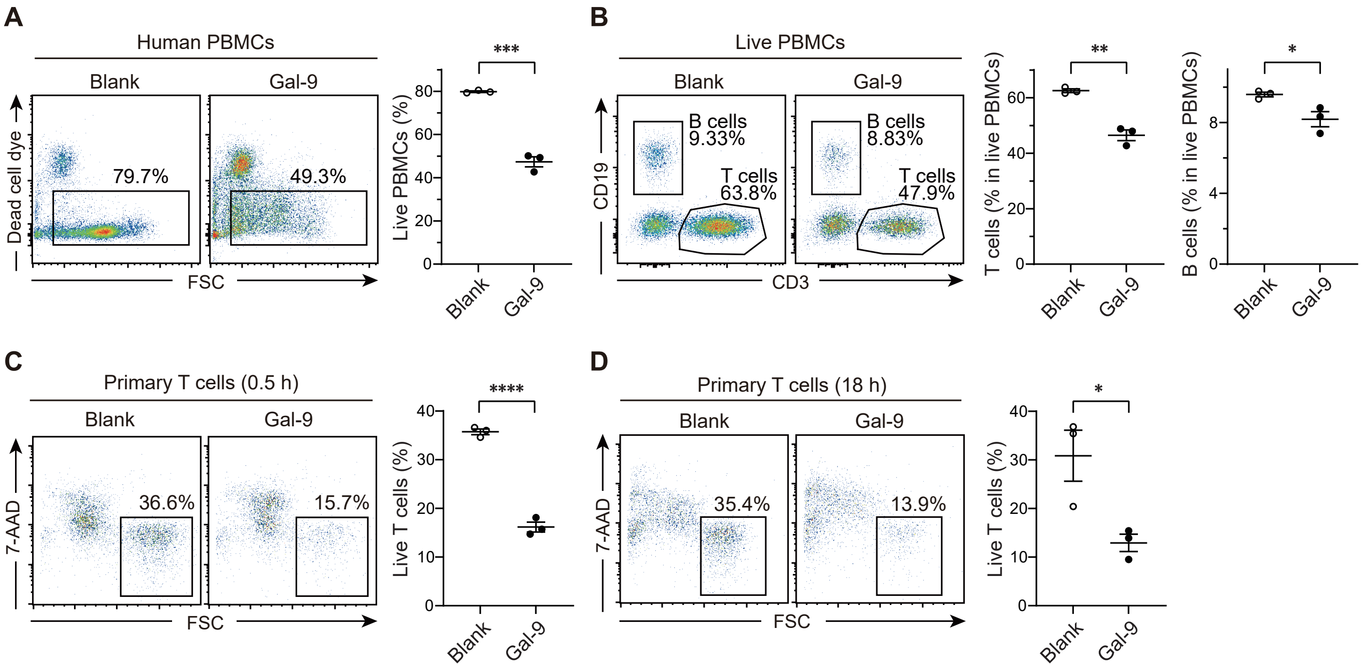
A
Human PBMCs
Blank
Gal-9
Dead cell dye
FSC
79.7%
49.3%
Live PBMCs (%)
***
Blank
Gal-9
B
Live PBMCs
Blank
Gal-9
CD19
CD3
B cells 9.33%
T cells 63.8%
B cells 8.83%
T cells 47.9%
T cells (% in live PBMCs)
**
B cells (% in live PBMCs)
*
C
Primary T cells (0.5 h)
Blank
Gal-9
7-AAD
FSC
36.6%
15.7%
Live T cells (%)
****
D
Primary T cells (18 h)
Blank
Gal-9
7-AAD
FSC
35.4%
13.9%
Live T cells (%)
*

Fig. S4

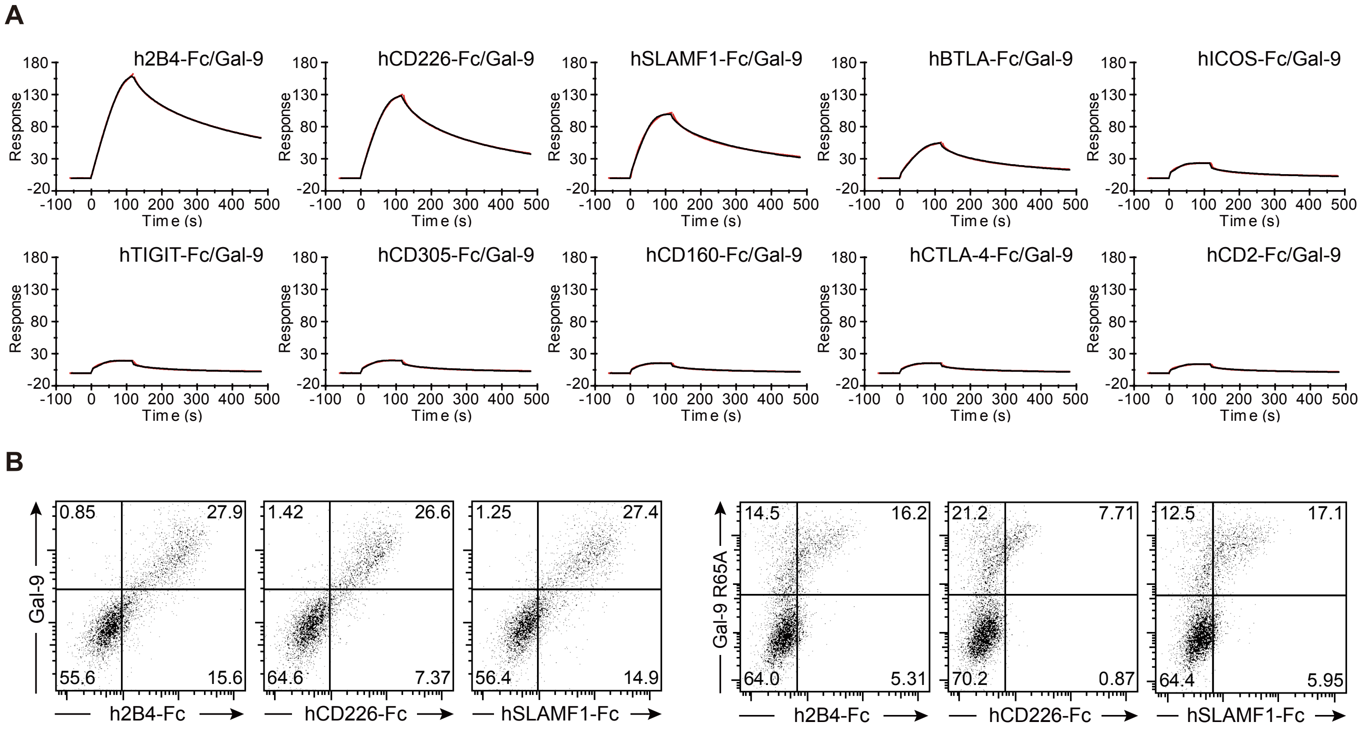
A
h2B4-Fc/Gal-9
hCD226-Fc/Gal-9
hSLAMF1-Fc/Gal-9
hBTLA-Fc/Gal-9
hICOS-Fc/Gal-9
hTIGIT-Fc/Gal-9
hCD305-Fc/Gal-9
hCD160-Fc/Gal-9
hCTLA-4-Fc/Gal-9
hCD2-Fc/Gal-9
Response
Time (s)
B
Gal-9
Gal-9 R65A
h2B4-Fc
hCD226-Fc
hSLAMF1-Fc
0.85
27.9
55.6
15.6
1.42
26.6
64.6
7.37
1.25
27.4
56.4
14.9
14.5
16.2
64.0
5.31
21.2
7.71
70.2
0.87
12.5
17.1
64.4
5.95

### Fig. S5

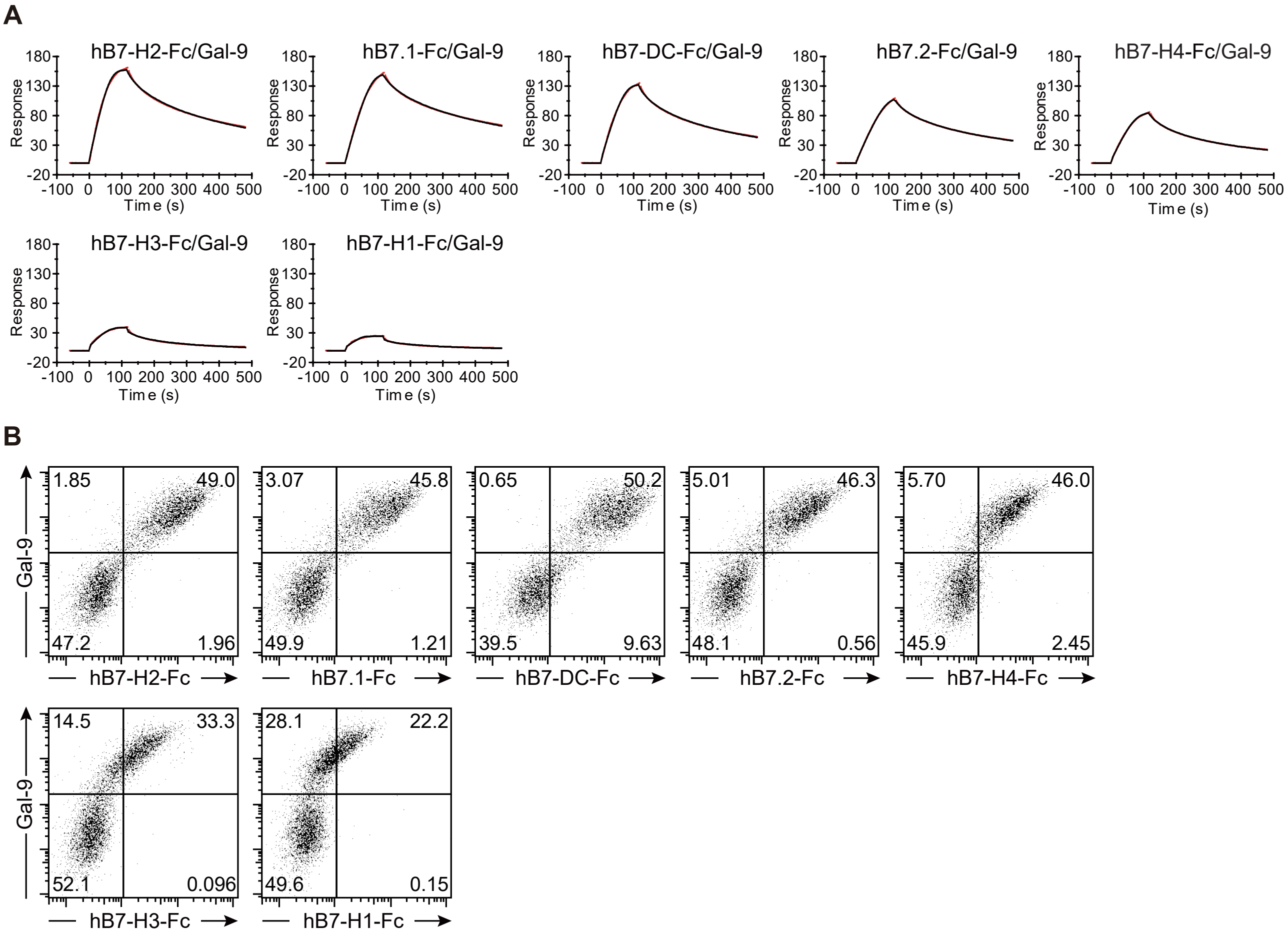
A
hBB7-H2-Fc/Gal-9
hB7.1-Fc/Gal-9
hB7-DC-Fc/Gal-9
hB7.2-Fc/Gal-9
hB7-H4-Fc/Gal-9
hB7-H3-Fc/Gal-9
hB7-H1-Fc/Gal-9
Response
Time (s)
B
Gal-9
1.85
49.0
47.2
1.96
hB7-H2-Fc
3.07
45.8
49.9
1.21
hB7.1-Fc
0.65
50.2
39.5
9.63
hB7-DC-Fc
5.01
46.3
48.1
0.56
hB7.2-Fc
5.70
46.0
45.9
2.45
hB7-H4-Fc
14.5
33.3
52.1
0.096
hB7-H3-Fc
28.1
22.2
49.6
0.15
hB7-H1-Fc

### Fig. S6

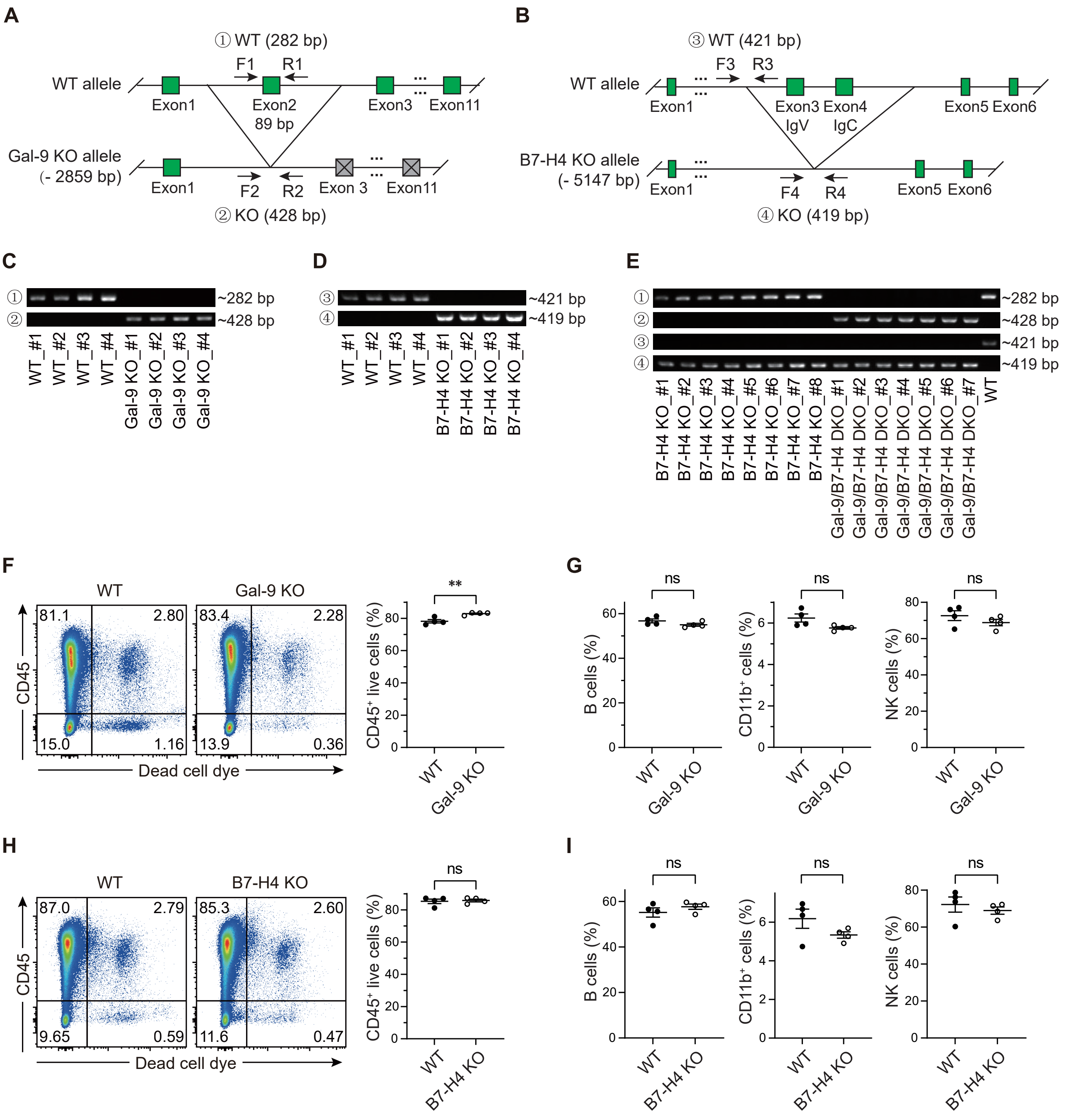

A
① WT (282 bp)
F1
R1
WT allele
Exon1
Exon2
89 bp
Exon3
Exon11
Gal-9 KO allele
(- 2859 bp)
F2
R2
Exon 3
② KO (428 bp)
B
③ WT (421 bp)
F3
R3
WT allele
Exon1
Exon3
Exon4
IgV
IgC
Exon5
Exon6
B7-H4 KO allele
(- 5147 bp)
F4
R4
④ KO (419 bp)
C
~282 bp
~428 bp
WT_#1
WT_#2
WT_#3
WT_#4
Gal-9 KO_#1
Gal-9 KO_#2
Gal-9 KO_#3
Gal-9 KO_#4
D
~421 bp
~419 bp
B7-H4 KO_#1
B7-H4 KO_#2
B7-H4 KO_#3
B7-H4 KO_#4
E
B7-H4 KO_#5
B7-H4 KO_#6
B7-H4 KO_#7
B7-H4 KO_#8
Gal-9/B7-H4 DKO_#1
Gal-9/B7-H4 DKO_#2
Gal-9/B7-H4 DKO_#3
Gal-9/B7-H4 DKO_#4
Gal-9/B7-H4 DKO_#5
Gal-9/B7-H4 DKO_#6
Gal-9/B7-H4 DKO_#7
WT
F
WT
Gal-9 KO
81.1
2.80
15.0
1.16
83.4
2.28
13.9
0.36
CD45
Dead cell dye
CD45⁺ live cells (%)
**
G
ns
B cells (%)
CD11b⁺ cells (%)
NK cells (%)
H
B7-H4 KO
87.0
2.79
9.65
0.59
85.3
2.60
11.6
0.47
I

Fig. S7

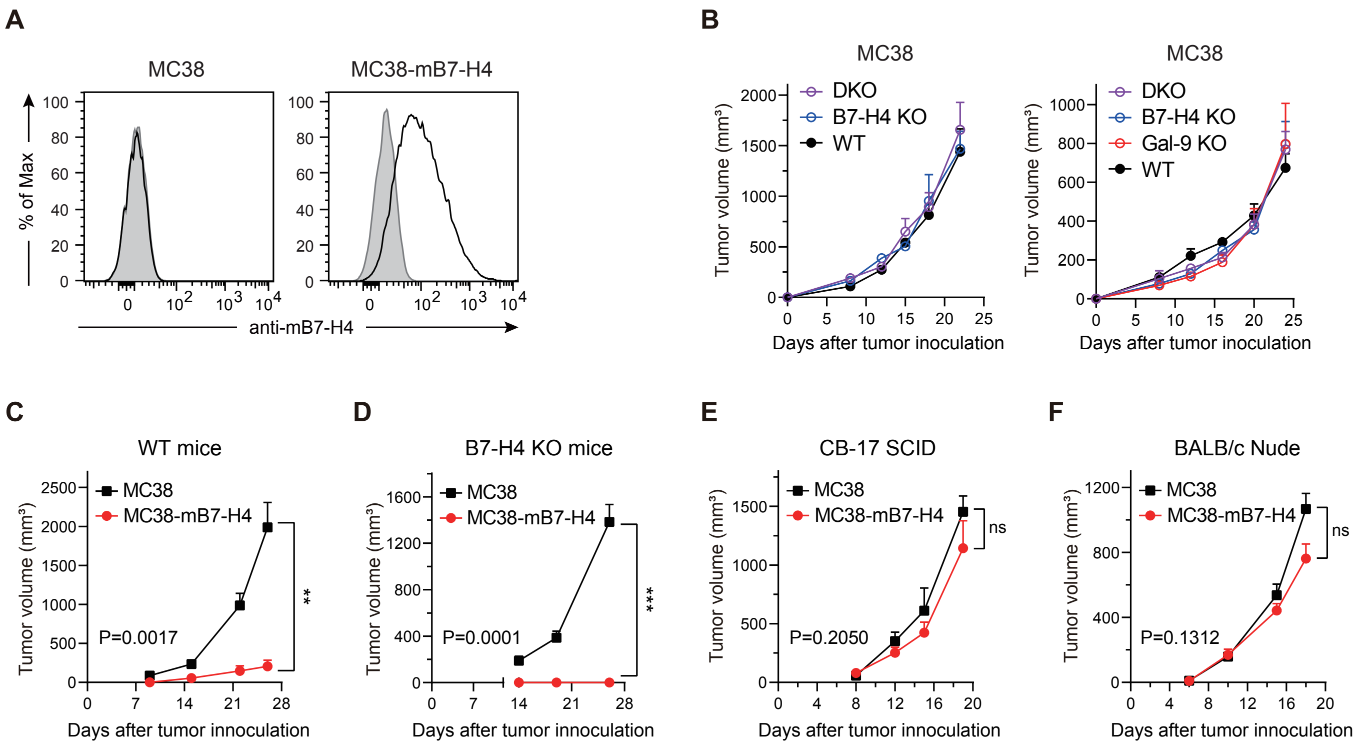
A
MC38
MC38-mB7-H4
% of Max
anti-mB7-H4
B
MC38
DKO
B7-H4 KO
WT
Gal-9 KO
Tumor volume (mm³)
Days after tumor inoculation
C
WT mice
MC38
MC38-mB7-H4
P=0.0017
**
D
B7-H4 KO mice
P=0.0001
***
E
CB-17 SCID
P=0.2050
ns
F
BALB/c Nude
P=0.1312
ns

**Table S1. Other reagents used in this article.**

| Product | Source | Catalog |
|---|---|---|
| Goat anti-Human IgG Fc Antibody, PE | Thermo Fisher Scientific | 12-4998-82 |
| Fixation/Permeabilization Kit | BD Biosciences | 554714 |
| Donkey anti-Mouse IgG (H+L) Secondary Antibody, Alexa Fluor™ 647 | Thermo Fisher Scientific | A-31571 |
| Tumor Dissociation Kit (mouse) | Miltenyi Biotec | 130-096-730 |
| DNase I | Merck | 10104159001 |
| 96 Well Solid White Flat Bottom Microplates | Corning | 3917 |
| Cell Line Nucleofector™ Kit V | LONZA | VCA-1003 |
| Glo Lysis Buffer, 1X | Promega | E2661 |
| Bright-Glo™ Luciferase Assay System | Promega | E2620 |
| rProtein A Beads 4FF | Smart-Lifesciences | SA015025 |
| PNGase F | New England Biolabs | P0704S |
| Dynabeads™ M-280 Streptavidin | Thermo Fisher Scientific | 11205D |
| ProteoSilver Stain Kit | Merck | PROTSIL2-1KT |
| PMSF | Beyotime | ST506 |
| SuperSignal™ Chemiluminescent Substrate | Thermo Fisher Scientific | 34580 |
| TRIzol Reagent | Invitrogen | 15596026 |
| PrimeScript™ RT Master Mix | Takara Bio Inc. | RR036a |
| KOD -Plus- | Toyobo | F0934K |
| T4 DNA Ligase | New England Biolabs | M0202S |
| Plasmid mini-extraction kit | TIANGEN | DP103 |
| In-Fusion® HD Cloning Plus | Takara Bio Inc. | 638910 |
